# Multi-omics of Circular RNAs and Their Responses to Hormones in Moso Bamboo (*Phyllostachys edulis*)

**DOI:** 10.1101/2022.10.23.513435

**Authors:** Yongsheng Wang, Huihui Wang, Huiyuan Wang, Ruifan Zhou, Ji Wu, Zekun Zhang, Yandong Jin, Tao Li, Markus V. Kohnen, Xuqing Liu, Wentao Wei, Kai Chen, Yubang Gao, Jiazhi Ding, Hangxiao Zhang, Bo Liu, Chentao Lin, Lianfeng Gu

**Affiliations:** College of Forestry, Basic Forestry and Proteomics Research Center, Fujian Provincial Key Laboratory of Haixia Applied Plant Systems Biology, Fujian Agriculture and Forestry University, Fuzhou 350002, China; College of Forestry, Fujian Agriculture and Forestry University, Fuzhou 350002, China; Department of Molecular, Cell and Developmental Biology, University of California, Los Angeles, CA 90095, USA

**Keywords:** Alternative splicing, circular RNAs, degradome, *Phyllostachys edulis*, phytohormones

## Abstract

Circular RNAs are endogenous non-coding RNAs with covalently closed structures, which have important functions in plants. However, their biogenesis, degradation, and function upon treatment with gibberellins (GA) and auxins (NAA) remain unknown. Here, we systematically identified and characterized expression patterns, evolutionary conservation, genomic features, and internal structures of circRNAs using RNase R-treated libraries from moso bamboo (*Phyllostachys edulis*) seedlings. Moreover, we investigated the biogenesis of circRNAs dependent on both cis- and trans-regulation. We determined details regarding the function of circRNAs, including their roles in regulating microRNA-related genes and modulating the alternative splicing of their linear counterparts. Importantly, we developed a customized degradome sequencing approach to detect microRNA-mediated cleavage of circRNAs. Finally, we present a comprehensive view of the participation of circRNAs in the regulation of hormone metabolism upon treatment of bamboo seedlings with gibberellins (GA) and auxins (NAA). Collectively, our study uncovers important features of circRNAs including overall characteristics, biogenesis, function, and microRNA-mediated degradation of circRNAs in moso bamboo.

## Introduction

In the time since the circular genomes of plant viroids [1] and hepatitis delta virus [2] were discovered, it has become clear that circular RNAs comprise a large class of covalently closed circular molecules that lack 5′ and 3′ ends. Intronic circular RNAs (ciRNAs) and exonic circular RNAs (circRNAs) are generated via intron lariat debranching and back-splicing of exons, respectively [3, 4]. Although the existence of circular transcripts has been known for at least 40 years [1], their importance has been likely underestimated due to their low abundance [5]. Recently, more intense investigation of such molecules has resulted from the ability to generate libraries enriched for circRNAs, and from the availability of circRNA-specific bioinformatics algorithms [6–8].

Back-splicing is much less completely understood compared to canonical splicing. Although most circular RNAs in mammals [3] and *Caenorhabditis elegans* [9] are processed from internal exons with long flanking introns containing inverted complementary sequences (ICSs) [3, 7, 10], formation of circular RNAs in other species such as *Drosophila melanogaster* [11] and *Oryza sativa* [12] does not require RNA pairing across flanking introns. In addition to such *cis*-regulation, recent studies have revealed that inhibiting RNA-binding proteins (RBPs) could directly alter circular RNA expression [13]. For instance, RBPs such as interleukin enhancer binding factor 3 of nuclear factor 90 isoform (NF90), interleukin enhancer binding factor 3 of 110 isoform (NF110), DExH-box helicase 9(DHX9), double-stranded RNA-specific adenosine deaminase (ADAR1), and KH Domain Containing RNA Binding (QKI) regulate the formation of exonic circular RNAs [9, 14–17]. Splicing factors such as FUS RNA binding protein (FUS), heterogeneous nuclear ribonucleoprotein L (HNRNPL) and QKI regulate circRNA expression by binding to the flanking introns of circRNAs [14, 18]. In addition, knock-down of core spliceosomal components such as SF3A and SF3B increase circRNA levels [13]. Thus, RPBs have important role in regulating biogenesis of circRNAs.

In addition to their biogenesis, the decay of circular RNAs directly influences their accumulation levels. Endonuclease RNase L initiates cleavage of circRNAs, and some circular RNAs containing microRNA-binding sites such as CDR1 antisense RNA (*CDR1as*) may be degraded by AGO2-mediated cleavage [19]. However, the mechanism of the degradation for most circRNAs remains elusive. Regarding circRNA function, both transcription and splicing of host genes can be modulated by specific circRNAs [4, 20]. For example, *CircSEP3* has been proposed to form an RNA:DNA hybrid (R-loop) with its cognate DNA locus in the nucleus; this R-loop alters exon skipping of linear SEP3, which influences floral homeotic phenotypes in *Arabidopsis* [21].

Overall, from existing data we remain far from understanding biogenesis, functions, and decay of circRNAs. In this study, we systematically identified and characterized circRNAs via RNase R-treated RNA-seq in moso bamboo seedlings, with or without gibberellin (GA) and auxin (NAA) treatments. Furthermore, we investigated whether circular RNAs might regulate the splicing of their corresponding linear RNAs due to R-loop structures. Importantly, we took advantage of a modified protocol for degradome-seq to detect several miRNA-mediated cleavage events. Finally, we determined that expression of circular RNAs could not only be modulated dynamically by GA or NAA hormones but also might affect several hormone-related genes in moso bamboo.

## Results

### Profile of circRNAs in moso bamboo seedlings

To enrich circRNAs in moso bamboo seedling samples, total RNAs from seedlings treated with double-distilled water (ddH_2_O), NAA, or GA were incubated with RNase R and Ribo-Zero. Sequences were then generated from three biological repeats (**Figure 1**A). In total, we detected 5105, 4461, and 7748 putative circular RNAs from ddH_2_O, NAA, and GA treatments, respectively (Figure 1B, Table S1). To independently test whether these sequences represent circular RNAs, we carried out PCR-based sequence validation with divergent primers for 8 randomly selected circular RNAs after RNase R treatment. This validation demonstrated that most circular RNAs were indeed resistant to exonuclease degradation. The exception was *circ-RAD16*, which was further discovered to include back-spliced junction sites by Sanger sequencing (Figure 1C). Thus, we conclude that *circ-RAD16* was bona fide circular RNA.

**Figure 1.**
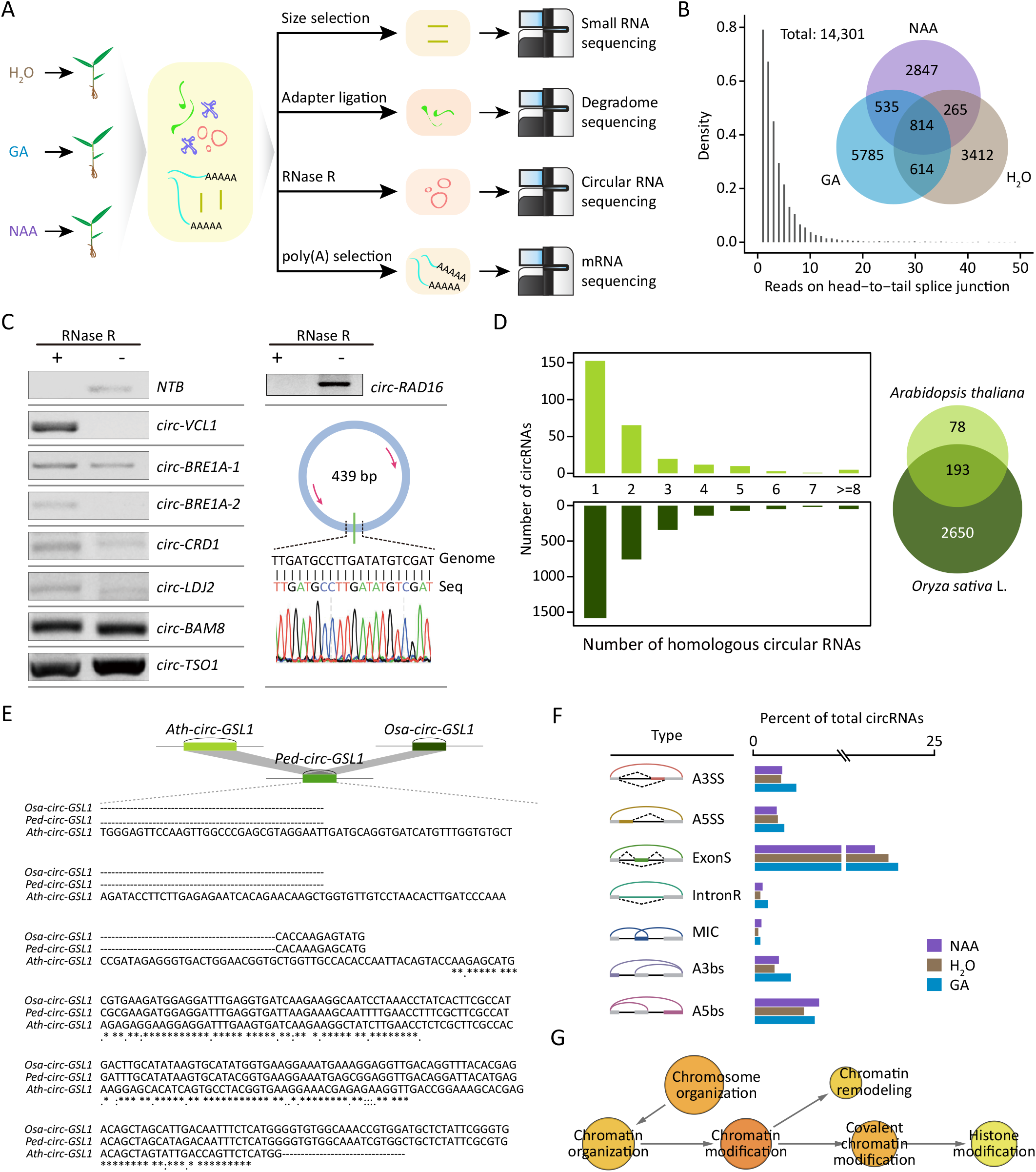
Characterization of circular RNAs in moso bamboo. **A.** Flow chart for multi-omics sequencing including mRNA sequencing, circular RNA sequencing, degradome sequencing, small RNA sequencing upon hormone treatment. **B.** Venn diagrams of the number of circular RNAs detected in different samples (upper panel). The density plot of back-splicing reads supported circular RNAs (lower panel). **C**. Validation of circular RNAs using RT-PCR after RNase R treatment with linear RNA (NTB) as control (left panel). Sanger sequencing validated back-splicing junctions of *circ-RAD16*, which was not resistant to RNase R (right panel). **D**. Number (left) and overlapping (right) of circular RNAs in *Arabidopsis* and *Oryza sativa* that show homology to circular RNAs in moso bamboo. **E**. Multiple-sequence alignment of conserved circ-*GSL1*. Asterisk symbols indicate highly conservative nucleotides. **F**. Diagrams of different AS types (left panel) and percentage of these circular RNAs (right panel). Gray bars and black lines represent exon and introns, respectively. Dotted curves and colored bars indicated AS events. Colored arclines represent back-splicing (circularization). **G**. GO enrichment analysis of circular RNAs with AS events. For more significant p-values, the node color gets increasingly more orange. The circle size was proportional to the number of genes enriched in the terms. The arrows represent hierarchical relations between GO terms. AS, alternative splicing.

We identified conserved circular RNAs by comparing our data with published data for other species [22]. As shown in Figure 1D, many circular RNAs of moso bamboo were homologous to a circular RNA of *Arabidopsis thaliana* or *Oryza sativa*. As expected, circular RNAs in bamboo exhibited more homology to those of *Oryza sativa* (2843, 19.8% of all circular RNAs) than to *Arabidopsis* (271, 1.8% of all circular RNAs). Notably, 193 (1.3% of all circular RNAs) were simultaneously detected in all three species; these included *circ-GSL1* (Figure 1E) generated from the gene encoding CALLOSE SYNTHASE 1 [23]. Enriched GO terms for these evolutionarily conserved circular RNAs included rhythmic processes, transporter activity and protein-containing complexes (Figure S1).

The complexity and diversity of circular RNAs are further increased by alternative back-splicing [6, 7, 24], which includes alternative 3’ splice site (A3SS), alternative 5’ splice site (A5SS), exon skipping (ExonS), intron retention (IntronR), alternative 3’ back-splice site (a3bs) and alternative 5’ back-splice site (a5bs) as well as mutually inclusive circular RNAs (MIC). CircRNAs generated by these seven types of alternative back-splicing were identified from all samples by circexplorer2 [7] (Figure 1F). Among the four AS types (A3SS, A5SS, ExonS and IntronR) that generated the different circRNA transcripts with same back-splicing sites, IntronR is the most prevalent in linear transcripts of plants, whereas ExonS was the most prevalent from the interior of circular RNAs. The other three types (a3bs, a5bs and MIC) generated different circRNAs with different back-splicing sites; a5bs accounted for the largest number of AS events among these three types. Enriched GO terms for these types of circular RNAs were involved in diverse biological functions, such as chromosome organization, chromatin remodeling and histone modification (Figure 1G).

### The biogenesis of circular RNAs is influenced by cis- and trans-regulation

Although circRNA-producing loci in moso bamboo usually had longer flanking introns than control introns (Figure S2), the circular RNAs themselves lacked obvious flanking intronic pairing sequences (**Figure 2**A), consistent with previous studies [3]. However, circRNA production can be driven by long artificial flanking inverted complementary introns [12]. To test for correlation of inverted complementary sequences with the biogenesis of circular RNAs, we cloned a 139-bp inverted complementary flanking intron with a representative exon from *CSLA1*, as a representative circRNA, and from the third linear exon of *NRT1*, which does not give rise to circular RNAs (Figure 2B). Semi-quantitative RT-PCR revealed that both the circularized exon and the linear exon in the overexpression vectors exhibited much higher circularization efficiency than in wild type, suggesting that RNA pairing with inverted complementary sequences could enhance back-splicing efficiency of circular RNAs. In particular, the linear exon from *NRT1* also could be induced into circular RNAs by inverted complementary sequences as flanking introns. To determine whether interior introns from multiple circularized exons could be involved in the biogenesis of circular RNA, we constructed *circ-PKL1* by including or excluding the interior intron. The presence of the interior intron did not lead to much difference in the rate of circRNA production (Figure 2C), suggesting that interior introns might not be key regulatory elements for the biogenesis of *circ-PKL1*.

**Figure 2.**
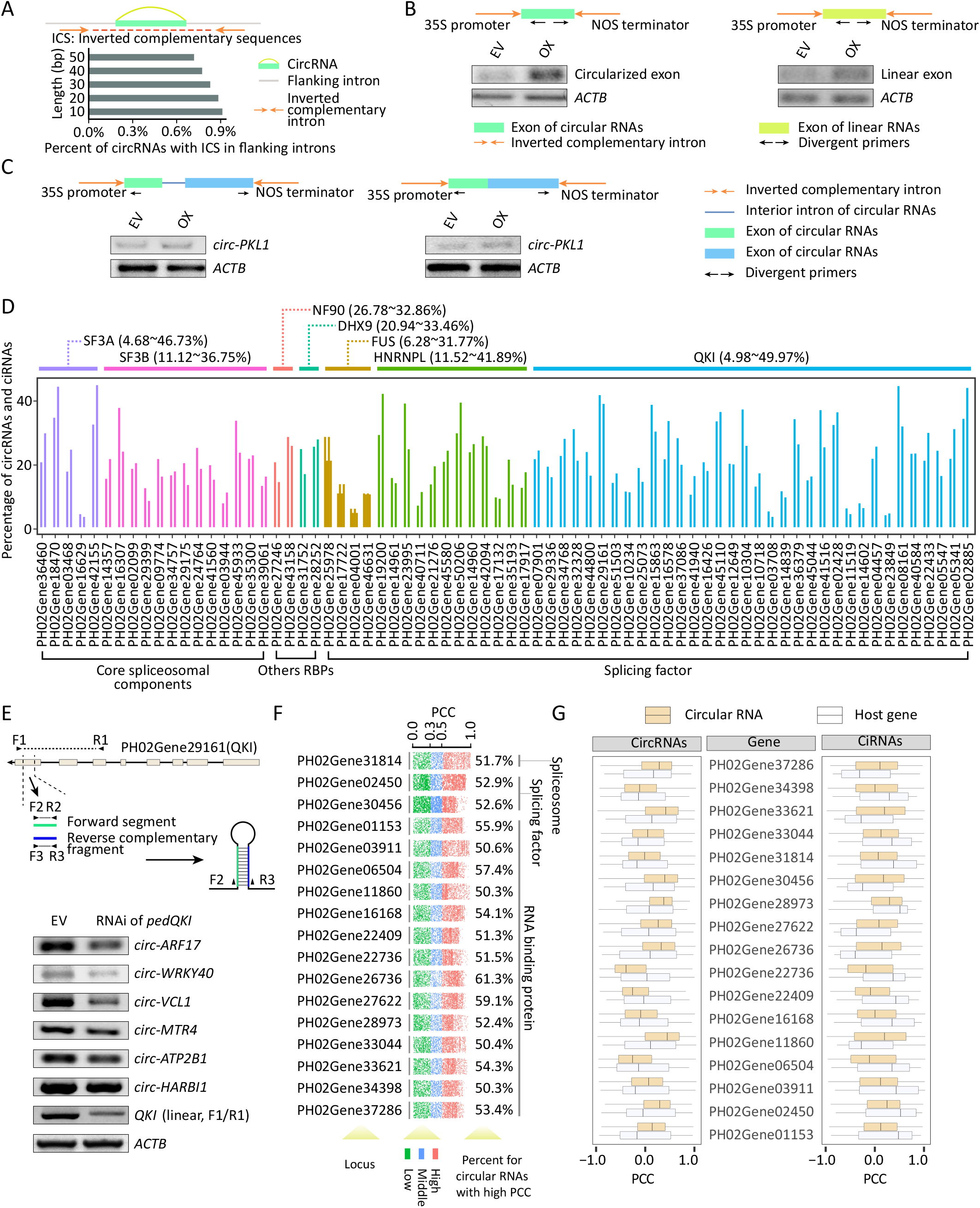
Biogenesis of circular RNAs in moso bamboo. **A**. Distribution of inverted complementary sequences (brown arrow) in flanking introns of circRNAs. Circularized exon and flanking intron are indicated by green bar and gray line, respectively. **B**. Schematic drawings show expression vectors including flanking inverted complementary sequences (brown arrow), circularized exon (green bar) and linear exon (yellow bar). PCR primers for circular RNAs are indicated as black arrows. Semi-quantitative RT-PCR in the lower panel shows circularization efficiency for circularized exon and linear exon, respectively. EV, and OX represent empty vector and overexpression vector, respectively. **C**. Experimental detection of the function for intron on multiple exon circularization of *circ-PKL1*. Schematic drawings of expression vectors including or excluding interior intron. Semi-quantitative RT-PCR in the lower panel shows circularization efficiency of *circ-PKL1* with or without interior intron. **D**. Percentage of circular RNAs showing high PCC (r>=0.5 or r<=−0.5) with proteins SF3A, SF3B, FUS, HNRNPL, QKI, NF90, and DHX9, respectively. **E**. RNAi Knock-down of *PedQKI*. Schematic drawing in the upper panel shows RNAi expression vector to knock-down expression of *PedQKI*. The lower panel shows PCR-based validation of the expression of six randomly selected circRNAs in QKI RNAi sample. **F**. PCCs between circular RNAs and 17 proteins including 1, 2, and 14 core spliceosomal factors, splicing factors and other RBPs. Number in right panel indicates detail percent of circular RNA with high PCC to those proteins, respectively. **G**. Distribution of PCCs between circRNAs/ciRNAs and linear RNAs generated from the same host genes.

Core spliceosomal components, splicing factors and some RNA-binding proteins (RBPs) have been also reported to participate in circular RNA regulation [7, 15, 25, 26]. To further explore the interplay between circular RNAs and these proteins, we performed sequence similarity analysis and identified 124 putative core spliceosomal components, 92 splicing factors, and 1132 other RNA-binding proteins excluding splicing factors and core spliceosomal components in moso bamboo. We also performed RNA-seq to calculate accumulation levels for linear transcripts (Figure 1A). Pearson correlation coefficients (PCCs) were calculated based on levels of circular RNAs (read per million, RPM) from circular RNA sequencing and expression of RNA-binding protein genes from RNA-seq (fragments per kilobase of exon per million fragments mapped, FPKM). The levels of circular RNAs were correlated (either positively or negatively; r>=0.5 or r<=−0.5) with the transcript levels for 44, 57, and 497 of core spliceosomal components, splicing factors, and other RBPs, respectively (Figure S3A and 3B, Table S2).

We focused on analyzing seven RNA-binding proteins (SF3A, SF3B, NF90, DHX9, FUS, HNRNPL, and QKI) that have been found to modulate the production of circular RNAs in other species. SF3A and SF3B, as core spliceosomal components, were encoded by 5 and 13 homologous genes in bamboo, respectively, and showed correlation in levels with 4.68~46.73% and 11.12~36.75% of circRNAs, (Figure 2D). In mammals, NF90 and DHX9 contain dsRNA binding domains (dsRBDs) and facilitate circular RNA formation by directly binding inverted repeated Alus (IRAlus) [16, 17]. Although the Alu element is rare in moso bamboo, NF90 and DHX9 exhibited correlation with 26.78~32.86% and 20.94~33.46% of circular RNAs, respectively (Figure 2D). In addition, the levels of 6.28~31.77%, 11.52~41.89% and 4.98~49.97% of circular RNAs (Figure 2D) were strongly correlated with those of transcripts encoding 4, 12, and 34 proteins homologous to splicing factors FUS, HNRNPL, and QKI, respectively. A striking example was PedQKI (PH02Gene29161) (Figure S4), which encoded a conserved KH domain and showed negative or positive correlation with 579 exonic and 423 intronic circular RNAs (a total of 49.97% of selected circular RNAs), respectively. Although expression of *circ-HARBI1* and *circ-ATP2B1* was not markedly altered, knock-down of PedQKI decreased the abundance of *circ-ARF17*, *circ-WRKY40*, *circ-VCL1* and *circ-MTR4* (Figure 2E).

Notably, 17 previously unreported proteins, namely 1 core spliceosomal factor, 2 splicing factors, and 14 other RBPs, were more highly correlated with circular RNAs than were the above seven RBPs (Figure 2F). We further explored the correlation between the expression of these 17 RBPs and of the host genes that generated these circular RNAs. We found that circular RNAs and their host genes exhibited distinct correlation patterns (Figure 2G). Taken together, these results suggest that core spliceosomal factors, splicing factors, and other RBPs might serve as regulators of the biogenesis of abundant circular RNAs. However, this analysis we identified potential RBP regulators by measure of linear dependence/correlation relationships using PCCs between the expression of circular RNAs and genes for RBPs. Thus, experimental analysis of RBP–RNA interaction will be required in the future to identify direct regulators of specific circRNA biogenesis by these RBPs.

### CircRNAs are involved in regulation of alternative splicing by R-loop structures

Our previous study revealed that overexpressing circ-*IRX7* in *Populus trichocarpa* decreased the rate of intron retention in the linear *IRX7* counterpart [27]. In this study, we used our RNA-seq and published data [28, 29] to identify AS events in bamboo, *Arabidopsis*, and *Oryza sativa*. We subsequently detected the locations of all AS events within circRNAs and ciRNAs across the three species (**Figure 3**A, 3B, and Table S3). To further validate if AS preferentially occurred within circular RNAs, we randomly extracted the same number of mRNA segments without producing circular RNAs and identified locations of AS events within these segments. The AS within the simulated random RNA segments had significantly lower frequencies than observed within circRNAs (Figure 3C). This trend was also observed in *Arabidopsis* and *Oryza sativa* (Figure 3C), which demonstrated that the elevated frequency of AS events within circRNAs might be conserved in different species. However, AS events located in the ciRNAs did not exhibit this trend (Figure 3D).

**Figure 3.**
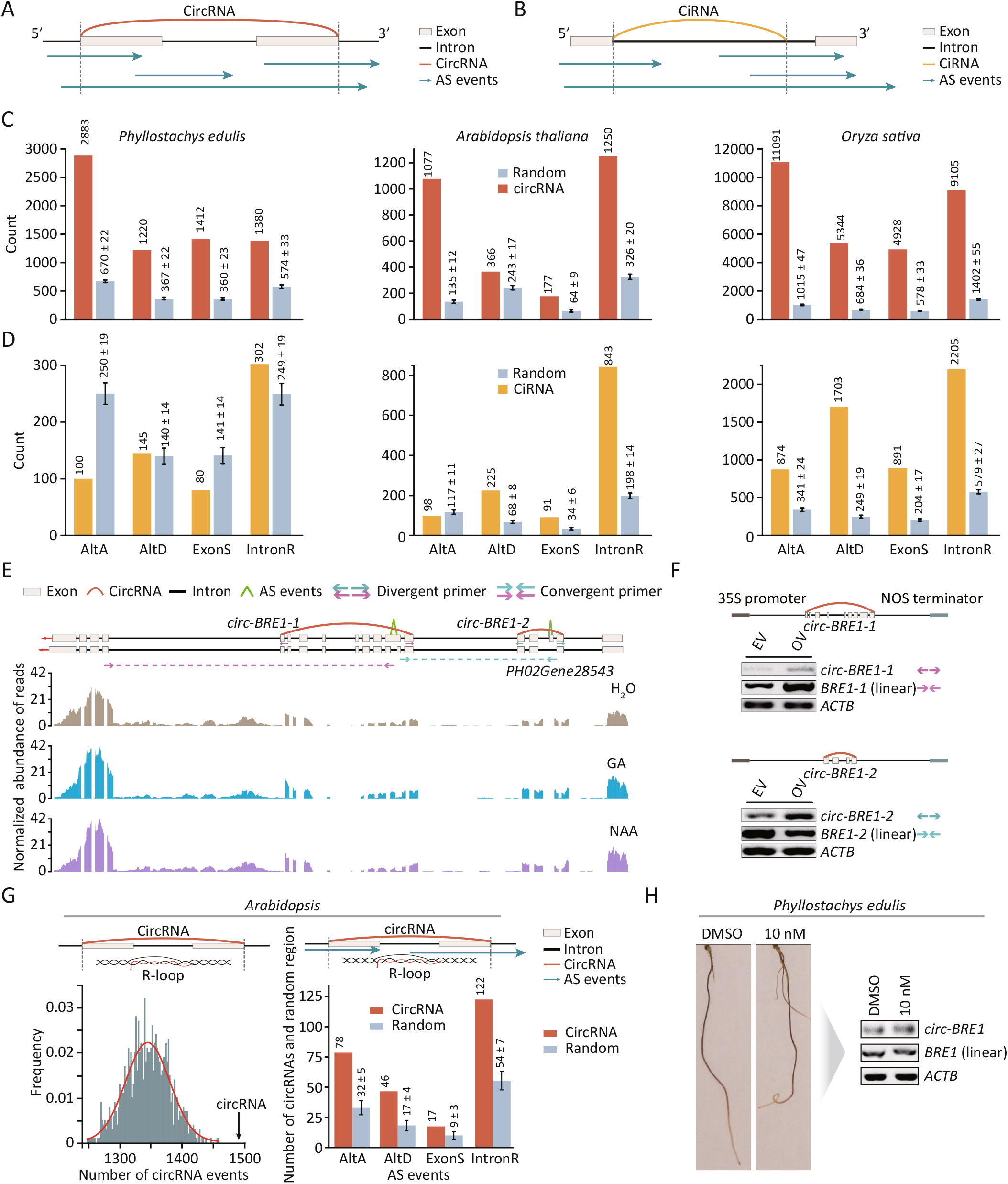
CircRNAs regulate alternative splicing in moso bamboo. **A-B**. Overview of overlapped regions between AS events and circular RNAs including circRNAs (A) and ciRNAs (B). Pale white bars indicate exons, black bars indicate introns, blue arrows indicate AS events, and colored arclines indicate back-splicing junctions. **C-D**. The histogram shows the number of AS events in the transcribed region of circular RNAs and random RNA segments in three species. Light yellow and brown bars represent circular RNAs (C) and ciRNAs (D), respectively. Steel gray bars represent random RNA segments. **E-F**. Visualization and validation of circRNAs such as *circ-BRE1-1* and *circ-BRE1-2* and their corresponding AS events. **E**. Visualization of intron retention and exon skipping events in transcribed regions of *circ-BRE1* and *circ-BRE2*, respectively. **F**. RT-PCR validation of *circ-BRE1-1* and *circ-BRE1-2* and their corresponding AS events, *linear-BRE1-1* and *linear-BRE1-2*. Divergent arrows represent divergent primers and convergent arrows represent convergent primers. Linear RNA of *actin* was used as a control. **G**. The histogram plot shows the overlap between R-loop and AS events in transcribed regions of circRNAs. Black arrow in the left panel indicates the observed number of circRNAs located in R-loop regions. **H**. Root phenotype upon DMSO and CPT treatment (left panel). Semi-quantitative RT-PCR shows the expression of *circ-BRE1-1* and *linear BRE1-1* upon CPT treatment (right panel).

PCCs between circRNAs and four types of AS events were calculated using their expression profiles according to circular RNA sequencing (RPM) and RNA sequencing (normalized reads that span splicing junctions), respectively. Approximately 19%~48% of circular transcripts in three species showed positive relationships with AS events (Table S3). To further validate the regulation of splicing by circRNAs, we selected *circ-BRE1-1* and *circ-BRE1-2*, which overlapped with ExonS and IntronR events from *E3 ubiquitin-protein ligase BRE1-like 1*, respectively (Figure 3E). Indeed, overexpression of *circ-BRE1-1* and *circ-BRE1-2* significantly changed the number of events in which the long isoforms of IntronR (*linear-BRE1-1*) and ExonS (*linear-BRE1-2*) were formed (Figure 3F), which is supportive of circular RNAs being able to modulate the AS events of their linear counterpart.

It has been shown that the presence of *circSEP3*, derived from an exon of *SEP3*, could result in exon skipping of its linear transcripts via an R-loop structure [21]. We therefore identified R-loop structures including AS events in transcribed regions of circular RNAs by integrating circular RNA, RNA-seq [29] and R-loop data in [30] *Arabidopsis*. Notably, transcribed regions of circRNAs had a significantly greater frequency of R-loop events compared to random regions (Figure 3G). Moreover, R-loop events were significantly more enriched in AS events than in random regions (Figure 3G). CPT is a TOP1 inhibitor that promotes R-loop accumulation in *Arabidopsis* [31]. We treated seedlings with DMSO or CPT (10 nm), which revealed that seedlings treated with CPT had enhanced abundance of *circ-BRF1* and long isoforms of IntronR (*linear-BRE1-1*) in transcribed regions of *circ-BRF1-1* (Figure 3H). Collectively, these findings indicated that circRNAs could regulate the AS frequency of their precursor transcripts, which may be related to R-loop structure formation by circRNA:DNA hybrids.

### Circular RNAs are involved in regulation of microRNA/siRNA-related genes

*CircAGO2* is generated from *AGO2* and represses AGO2/miRNA-mediated gene silencing [32]. In total, 163 miRNA-related genes were identified by sequence similarity analysis, and 79 miRNA-related genes including AGO1 and SDN1 were identified as host genes of circular RNAs in moso bamboo (**Figure 4**A and Table S4). We selected *circ-DCL4*, which originates from *DCL4*, for further validation. An RNase R exonuclease experiment revealed the resistance to exonuclease, and sequencing also validated the back-splice sites in *circ-DCL4* (Figure 4B). Notably, overexpressing *circ-DCL4* tended to decrease the expression level of its linear RNA (*linear-DCL4*) (Figure 4C), which might explain its effect on miRNAs/siRNA biogenesis.

**Figure 4.**
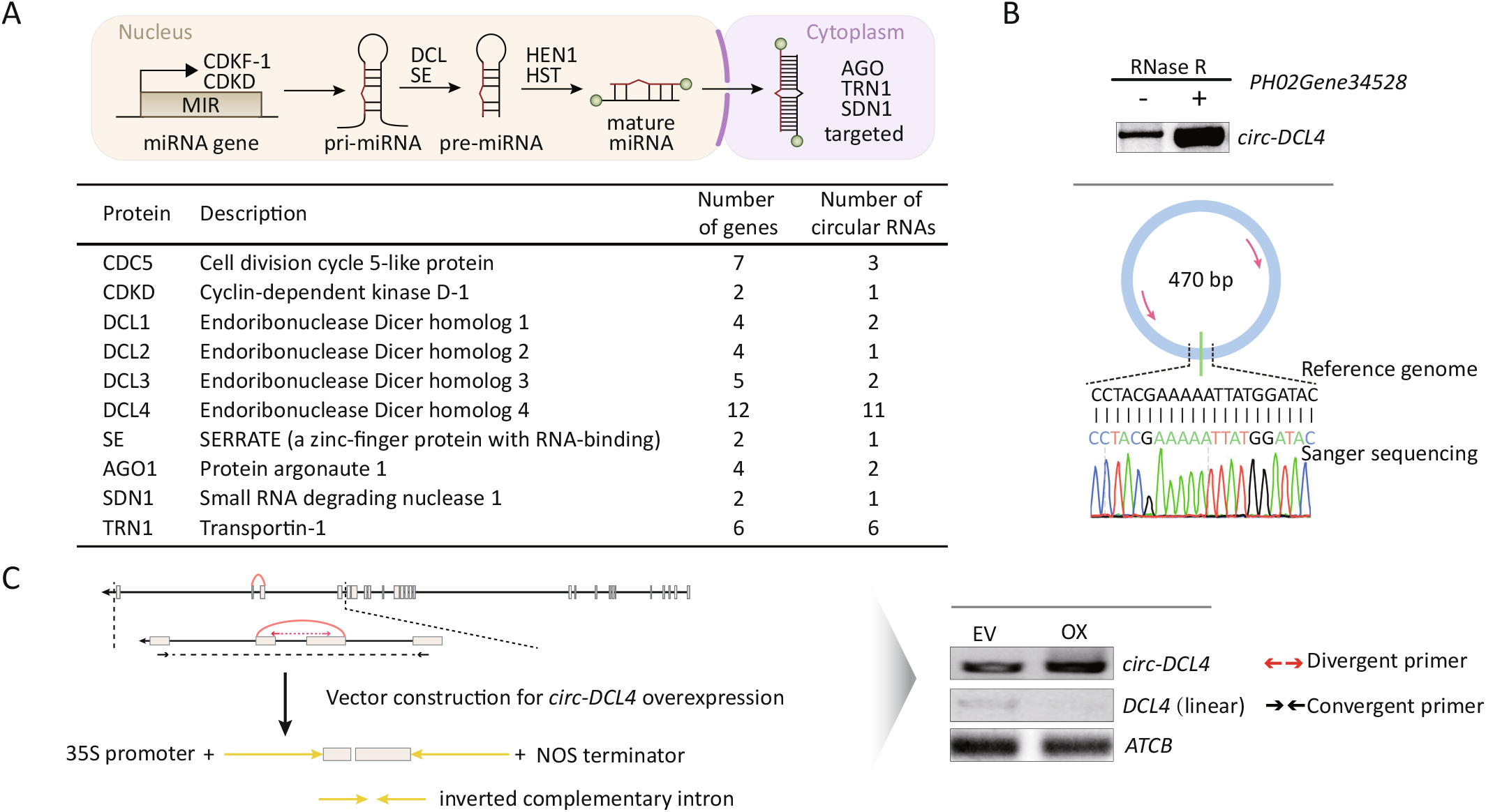
Circular RNAs regulate miRNA-associated genes in moso bamboo. **A**. The upper panel shows the miRNA-associated genes involved in several major steps in miRNA biogenesis and modes of action in plants. The table in the lower panel shows number of miRNA-associated genes and their corresponding circular RNAs. **B**. RT-PCR and Sanger sequencing for validation of *circ-DCL4* with RNase R treatment. **C**. The left panel shows the vector construction for *circ-DCL4* and *linear-DCL4*. RT-PCR validation in the right panel shows the overexpression of *circ-DCL4* and expression of *linear-DCL4*.

### Circular RNAs are translatable in moso bamboo

In this study, we developed a computational pipeline for detecting translatable open reading frames of circRNAs (cORF) using proteomics (Figure S5A). Moreover, we identified upstream ORF (uORF) and downstream (dORF) linear transcripts. Annotated CDS regions for each gene were regarded as primary ORFs (pORFs). As shown in Figure S5B, a small proportion of cORFs, uORFs, and dORFs showed sequence similarty to entries in Nr, PsORF, and UniProt. For instance, a conserved cORF derived from *circ-GLO5* was homologous to small ORFs in plants from the PsORF database [33], which indicated that these circRNAs might encode functional peptides (Figure S5C). As the third step, all ORFs were used as a search library for LC-MS/MS–based proteomics to identify unique peptide evidence for each ORF. In total, 538 cORFs from 536 circRNAs (approximately 7.1% of all 7554 circRNAs) were predicted to be translatable based on proteomics evidence (Figure S5D, Table S5). For example, *circ-P4H-1*, generated a detectable protein with a length of 289 aa spanning the back-splicing site with evidence of a unique peptide (Figure S5E).

### MiRNA-mediated cleavage of circRNAs

Endonucleases RNase L have been reported to decay circular RNAs globally in animals [34]. Sequence similarity analysis of RNase L from 67 animals revealed a high identity (on average 80.5%) and alignment length (on average 727 aa). By contrast, we did not identify Ribonuclease L sequences in 100 plants including moso bamboo (Figure S6A, S6B), suggesting that the decay of circular RNAs in plants is not mediated by RNase L. Given that miR-671 was identified as a key regulator in the decay of CDR1as [19], we hypothesized that miRNAs may also be potential factors contributing to the decay of circular RNAs in plants. We began by sequencing 9 small RNA libraries constructed from the same material as circular RNAs (Figure 1A). We detected 823 mature miRNAs including 164 conserved miRNAs and 659 variant miRNAs, which clustered to form 43 miRNA families (Figure S7A, Table S4). Conventional degradome sequencing [35] could not detect the cleavage of circular molecules due to the absence of 3◻ poly(A) tails of circular RNAs. Therefore, we modified the protocol for generating degradome libraries so that we could identify miRNA-mediated cleavage of circRNAs (**Figure 5**A). After poly(A) selection, total RNAs were separated into poly(A)− and poly(A)+ transcripts. Subsequently, the poly(A)− transcripts were subjected to rRNA depletion followed by immediate ligation to 5’ RNA adapters to enrich the cleavage transcripts of circular RNAs including free 5’ monophosphate, whereas the 5’ monophosphate of poly(A)+ transcripts was directly ligated to the 5’ RNA adapter. Finally, these two types of degradome libraries were prepared by reverse transcription and sequenced. As expected, transcript regions including 5◻ UTR, CDS and 3◻ UTR were more enriched with free 5’ monophosphate reads in the poly (A)+ library than in the poly (A)-library (Figure 5B). Consistent with previous observations that the cleavage transcripts were biased toward the 3’ end of mRNAs [36], poly(A)+ library also tended to generate more reads including free 5’ monophosphate in the 3’ UTR than in the 5’ UTR. However, the poly(A)− library did not exhibit the tendency (Figure S7B).

**Figure 5.**
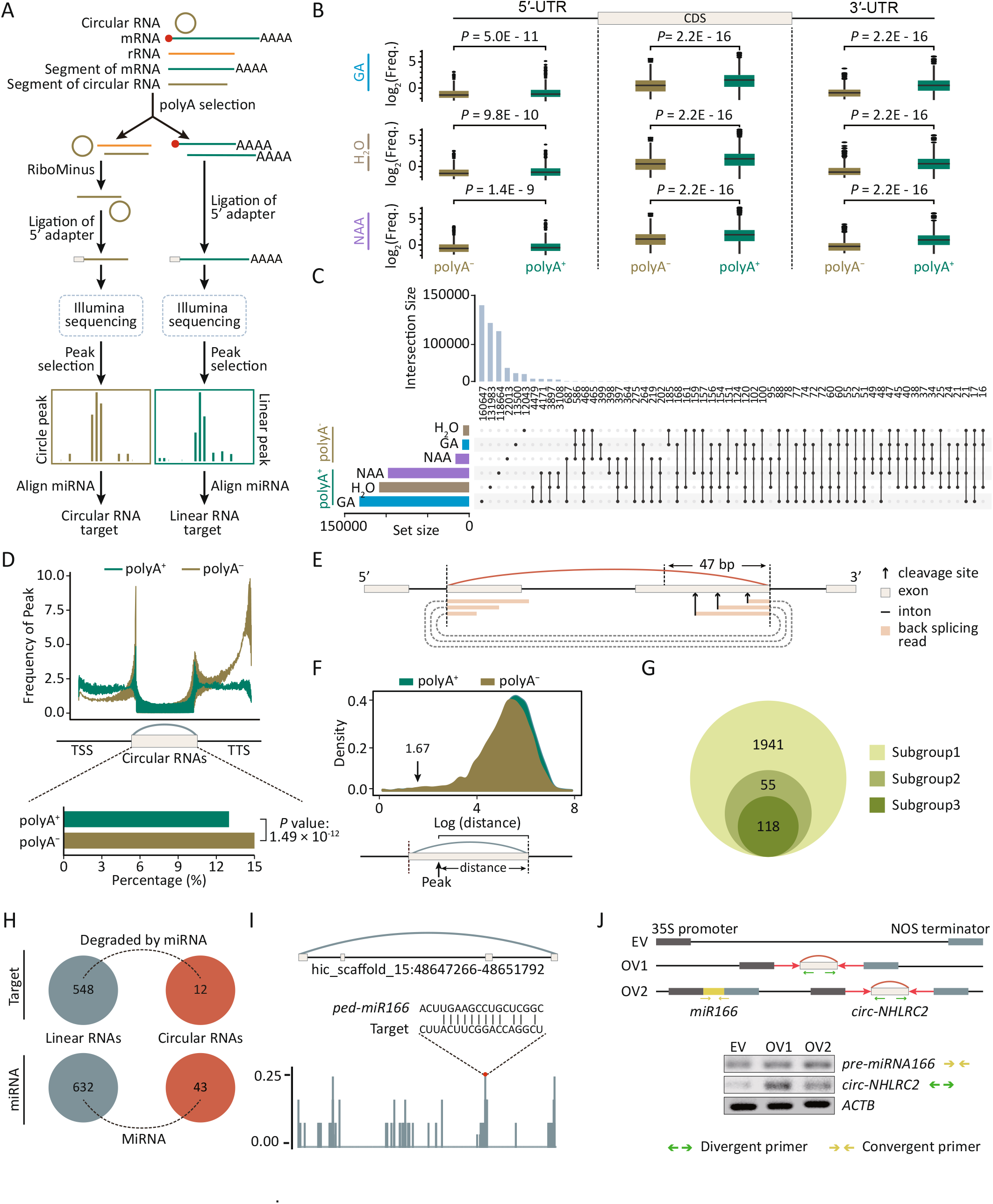
MiRNA-mediated cleavage of circular RNAs in moso bamboo. **A**. Flow chart for construction of customized degradome libraries for enriching the decaying circular RNA without poly(A) tails. **B**. The box line plot shows the distribution of reads in 5◻ UTR, CDS and 3◻ UTR of the gene from poly(A)+ and poly(A)− degradome libraries. **C**. The UpSet plot shows the intersection of cleavage sites among different types of libraries and different hormone treatment. The histogram plot in upper panel shows the number of cleavage sites shared by different libraries. The histogram plot in the lower panel and the line plot in the right panel show total cleavage sites from each library and the combination of different library. **D**. Plot in the upper panel shows the distribution of cleavage sites from two types of libraries in genes divided into three groups: the upstream regions of circular RNA body, downstream regions of circular RNA body and circular RNA body. Bar in lower panel shows the percentage of cleavage sites from two types of libraries in transcript regions of circular RNAs. **E**. Schematic overview of decaying reads of circular RNAs spanning back-splicing sites. Pale white bars represent exons, black lines represent introns, black arrows represent cleavage sites, and colored arclines represent back-splicing. Light yellow bars and dash arclines indicate the mapped back-splicing junction reads. **F**. Length distribution between cleavage sites and the 3’ end of transcribed regions of circular RNAs. **G**. Venn diagram shows the number of three subgroups. Subgroup 1, 2, 3 represents the number of circular RNAs including cleavage sites based on degradome sequencing, the number of circular RNAs with a distance less than 47 nucleotide between cleavage sites and 3’ end of transcriptional regions of circular RNAs, and the number of circular RNAs determined by degradome reads spanning back splicing sites. **H**. Number of circular RNAs and linear RNAs including miRNA-mediated cleavage. **I**. Schematic overview of miR156-mediated cleavage sites in *circ-NHLRC2*. **J**. The upper panel shows the vector construction for empty vector (EV), *circ-NHLRC2* (ov1) and plasmid expressing with both *circ-NHLRC2* and miRNA166 (ov2). The lower panel indicates RT-PCR detected the expression of *circ-NHLRC2* and miRNA166. Divergent and convergent arrows represent divergent and convergent primers, respectively.

We further developed a computational strategy to identify miRNA cleavage events in circular RNAs, termed ‘degradome peaks in poly(A)− transcripts’ (Figure S7C). In brief, degradome sequencing reads were first aligned to the genome using Bowtie2 [37] to detect accumulated cleavage events termed ‘degradome peaks’. As indicated in Figure 5C, degradome peaks within annotated transcript regions from each poly(A)+ library were considerably higher than those in the poly(A)− library from the same samples. The distribution of degradome peaks was significantly enriched in upstream and downstream regions of circular RNAs, regardless of whether the peaks originated from poly(A)+ or poly(A)− libraries (Figure 5D, upper panel). We further compared the distribution of degradome peaks between poly(A)− and poly(A)+ libraries in host genes that generated circular RNAs. We observed that the frequency of degradome peaks from the poly(A)− library (approximately 15%) was slightly higher (*P*-value=1.49e-12, Fisher test) than that from the poly(A)+ library (approximately 13%) in the region of circular RNAs (Figure 5D, lower panel).

We calculated the distance from peaks to back-splicing sites to search for the decaying peaks spanning back-splicing sites (Figure 5E and 5F). The degradome reads from these peaks were then aligned to the upstream and downstream 50 nucleotides of back-splicing sites (Figure S7D). In total, degradome peaks from 118 circular RNAs were revealed by degradome reads spanning back-splicing sites (Figure 5G). These results collectively suggest that our customized libraries and computational pipeline could effectively identify the decaying sites of poly(A)− transcripts, particularly for circular RNAs.

We further identified degradome peaks in the poly(A)− library originating from the miRNA-mediated cleavage of circular RNAs. The upstream and downstream 25 nucleotides of significant degradome peaks were aligned to mature miRNAs by RNAplex [38] to identify candidate miRNA cleavage sites in the ±1bp region of degradome peaks. Overall, 12 circular RNAs and 548 linear RNAs were identified as the cleavage transcripts mediated by 43 and 632 miRNAs, respectively (Figure 5H and Table S6). For instance, we observed that ped-miRNA166 mediated cleavage in *circ-NHLRC2* (Figure 5I). For further validation, we overexpressed *circ-NHLRC2* (OV1) and both *circ-NHLRC2*/miR166 (OV2) (Figure 5J). Semi-quantitative RT-PCR revealed that the expression of *circ-NHLRC2* was significantly increased in protoplasts transformed and overexpressing only *circ-NHLRC2* (OV1). By contrast, *circ-NHLRC2* exhibited a decreased tendency in the OV2 vector carrying both *circ-NHLRC2* and miR166 (Figure 5J). Taken together, these observations suggest that miRNAs could contribute to the degradation of circular RNAs in moso bamboo.

### CircRNA expression is modulated by GA and NAA

The plant hormones GA and NAA are essential for developmental processes in moso bamboo including root germination and shoot development [39, 40]. However, whether circular RNAs respond to hormones has remained unknown. To perform quantitative analyses of circular RNAs, we extracted back-splicing junctions relative to normalized circRNA abundance using RPM. Strikingly, the percentage of upregulated circular RNAs upon GA treatment was two-fold more than that of downregulated circular RNAs (27.1% upregulated and 12.9% downregulated), whereas the many fewer were upregulated than downregulated upon NAA treatment (3.9% upregulated and 22.3% downregulated) (**Figure 6**A and 6B, and Table S7).

**Figure 6.**
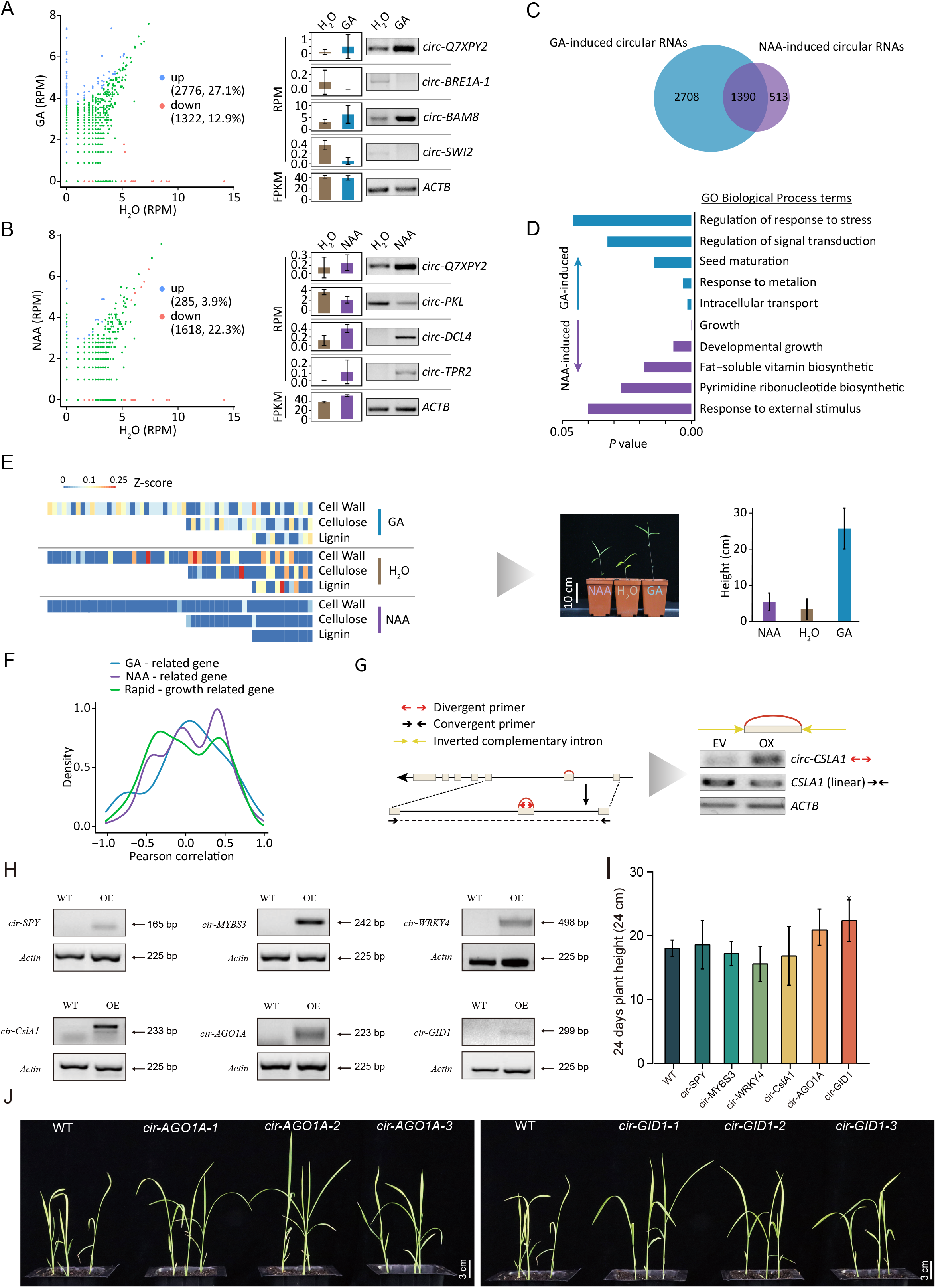
Hormone-induced circular RNAs in moso bamboo. **A**. The scatter plot in the left panel shows RPMs for H_2_O (x-axis) and GA treatments (y-axis). Semi-quantitative PCR using divergent primers in the right panel validated the differential expression data of circular RNA in response to GA treatments with linear transcripts actin as internal reference gene using convergent primers. **B**. The scatter plot shows the differential levels of circular RNA in response to NAA and semi-quantitative PCR shows the validation of the levels of circular RNAs based on sequencing. **C**. Venn diagram shows overlapped and unique differential circular RNAs in response to GA and NAA. **D**. The histogram plot shows the top 10 GO terms enriched for hormone-induced circular RNAs. **E**. Heatmap showing expression levels of circular transcripts related to cell wall, cellulose, and lignin. Red or blue represents high and low abundance of circular RNAs, respectively. Plot in lower panel shows phenotype and height of seedlings upon GA and NAA treatment after two-weeks. **F**. The PCCs of circular RNAs and their host linear RNAs related with fast-growth, NAA, and GA. **G**. The vector construction (left panel) and RT-PCR validation (right panel) of *circ-CSLA1* and *linear-CSLA1*. Divergent and convergent arrows represent divergent and convergent primers, respectively. **H**. RT-PCR using divergent primers revealed the expression level of circRNAs in six transformed lines (T1 generation). **I**. The histogram shows the plant height from 24-day-old seedlings. **J**. The phenotypes of plant height for *cir-SPY*, *cir-MYBS3*, *cir-WRKY4*, *cir-CslA1*, *cir-AGO1A*, and *cir-GID1*.

Furthermore, the expression patterns of 7 tested circular RNAs were found to be consistent with circular RNA-seq analysis using semi-quantitative RT-PCR (Figure 6A and 6B). For instance, *circ-BRE1-1* from *E3 ubiquitin-protein ligase BRE1-like 1* (PH02Gene28543) displayed much lower levels with GA hormone treatment whereas NAA treatment resulted in higher levels of *circ-DCL4*. These findings indicate that both GA and NAA treatments induced significant changes and that alteration of circRNA levels was more sensitive to GA than to NAA (Figure 6C). Furthermore, transcript levels of 1,390 circular RNAs were both modulated by GA and NAA, as exemplified by *circ-Q7XPY2* (Figure 6C). GO enrichment analysis for differentially expressed circular RNAs revealed that intracellular transport and response to external stimulus were highly enriched in response to GA and NAA treatment, respectively (Figure 6D).

To further investigate the biological effects of hormone-induced circular RNAs, we clustered the expression of circular RNAs from different samples. As indicated in Figure S8A, the host genes for 67 significantly differentially expressed circular RNAs, were annotated as being related to second messengers, cell inclusion, plant organs, and plant hormones, namely the abscisic acid and cytokinin pathways. Moreover, circular RNAs resulting from rapid growth–related genes, including those related to the cell wall, cellulose, and lignin, were modulated in response to GA and NAA, which is consistent with the phenotype of seedling growth after hormone treatment (Figure 6E).

The regulation of concentration gradients for gibberellin and auxin hormones is a key process in plants [41, 42]. In total, 15 GA-related and 87 NAA-related genes were identified as parent genes that could produce circular RNAs (Table S7). These circular RNAs were likely to mediate biogenesis, function and transport of gibberellins (Figure S8B) and auxins (Figure S8C). For example, *circ-CPS1* and *circ-PhYUC5* resulted from PH02Gene47426 (*CPS1*) coding ent-copalyl diphosphate synthase 1 and PH02Gene12216 (*PhYUC5*) coding FMO-like, which are involved in gibberellin and auxin biosynthesis, respectively. PCC analysis between circular RNAs (RPM) and their linear RNAs (FPKM) indicated that 89 circular RNAs correlated in levels with their corresponding linear RNAs (Figure 6F). A striking example was *circ-CSLA1*, which is derived from *Glucomannan 4-beta-mannosyltransferase 1* (*CSLA1*) and is involved in generating the backbone used for galactomannan synthesis by galactomannan galactosyltransferase [43]. *Circ-CSLA1* was upregulated in response to GA treatment. Overexpression of *circ-CSLA1* significantly reduced the levels of its linear RNA (Figure 6G). Collectively, our results indicated that alteration of circRNA expression in response to GA and NAA might also modulate the expression of host genes.

As stable transformation is difficult in moso bamboo and both bamboo and rice are Poaceae family members [44], we overexpressed 6 candidate circRNAs (*cir-SPY*, *cir-MYBS3*, *cir-WRKY4*, *cir-CslA1*, *cir-AGO1A*, and *cir-GID1*) in rice via Agrobacterium-mediated transformation. Stably transformed lines showed successful transgene expression of six circRNAs (Figure 6H). Among these six transformed lines, *cir-AGO1A* and *cir-GID1* promoted plant height in three independent lines (Figure 6I, J, and Figure S9), which highlighted that circRNAs can have biological roles in plant.

## Discussion

Techniques involving profiling with poly(A)− RNA populations or RNase R enrichment have uncovered global expression of circular RNAs [25]. In this study, our comprehensive sequencing of an RNase R-treated library identified a total of 14,301 circular RNAs (Figure 1A and 1B), suggesting that this type of library could enhance the identification of circular RNAs. These expanded circular RNAs raised the possibility of investigating the expression pattern, evolutionary conservation, and the internal structure of circular RNAs. Notably, the homologous circRNAs were present in low proportions (Figure 1D), which may be explained by the tissue-specific expression. Consistent with previous findings [6, 7, 15], our identified circular RNAs harbored alternative back-splicing (ABS), alternative splicing (AS) events and mutually inclusive circular RNAs (MIC) (Figure 1F). We also used different methods to detect circRNAs. In total, over 60% of genes with circRNA AS events from CIRI-AS [45] overlapped with those from circexplorer2 [7], which indicated that most of these events could be identified by the different methods.

Emerging studies have revealed that circular RNA formation is influenced by cis-regulatory elements such as ICSs flanking exons [3] and trans-regulatory factors [7, 15, 25, 26]. Our study strongly indicates that artificial RNA pairing with inverted complementary sequences could enhance back-splicing efficiency in moso bamboo (Figure 2B), although the exons of circular transcripts lack native flanking intronic pairing sequences (Figure 2A). Intriguingly, overexpression of circular RNAs with included or excluded interior introns suggests that interior introns contained within multiple circularized exons may not be a key regulatory element for the biogenesis of circRNAs (Figure 2C). However, an extended investigation of the splicing of interior introns in other circular RNAs is required to obtain a more comprehensive conclusion.

Several known trans factors reported to participate in circular RNA regulation [7, 15, 25, 26] also exhibited strong associations with circular molecules in our samples (Figure 2E). For example, QKI, was more closely correlated with 579 exonic circRNAs than with 423 intronic circular RNAs. Knock-down analysis further confirmed the alteration in abundance for four selected circRNAs (Figure 2E). However, we still do not know whether there is direct regulation of these RPBs on specific motifs or circRNA transcripts due to the lack of RBP–RNA interaction experiment. It will be interesting to investigate these trans-regulatory factors for binding motifs around circular RNAs using high-throughput sequencing of RNAs isolated by cross-linking immunoprecipitation (CLIP-seq), for instance [46].

Combining RNase R-treated circular RNA libraries with a common RNA-seq library provided us the opportunity to comprehensively explore the relationship between circular RNAs and the AS of their linear counterparts. We identified several potential regulatory circRNAs, such as *circ-BRE1-1* and *circ-BRE1-2*, which like *circSEP3* [21], affected the AS of their precursor transcripts in *Phyllostachys edulis*, *Arabidopsis*, and *Oryza sativa* (Figure 3C, 3D, 3E, and 3F). Subsequently, we observed that transcribed regions of circRNAs overlapped R-loop events and AS events with significantly greater frequency than random regions (Figure 3G). The abundance of long isoforms (*linear-BRF1*) was increased after treatment with TOP1 inhibitor, which functions in promoting R-loop accumulation. This result indicated that R-loop structure could be a potential AS regulator of host genes. Furthermore, *circ-BRF1* also exhibited the identical trend with *linear-BRF1*. However, further research is essential to elucidate the relationships between R-loops, AS, and circRNAs, for example using knock-down analysis of R-loop and ssDRIP-seq [30], which should reveal whether R-loop accumulation can influence circRNA abundance and thus promote AS at the genome-wide level.

Our data showed that partial circular RNAs could originate from miRNA-related genes. A striking sample was *circ-DCL4*, which was validated by RNase R exonuclease treatment and Sanger sequencing (Figure 4B). Overexpression of *circ-DCL4* reduced the expression levels of *linear-DCL4* significantly (Figure 4C), which suggested that circular transcripts might participate in miRNA biogenesis, activity and degradation by modulating the level of miRNA-associated genes. The correlation in levels between circular RNAs and 823 mature miRNAs provided evidence for this hypothesis (Supplemental Tables S4). In the future, improvement of Agrobacterium-mediated transformation efficiency in moso bamboo would allow work to identify how many miRNAs or siRNA are changed in expression upon overexpressing these specific circular RNAs.

We found that plants appear to lack homologs of human RNase L (Figure S6A and S6B), which led us to test whether miRNAs contribute to the initiation of degradation of circular RNAs, similar to linear RNAs in plants. Degradome sequencing is the gold standard for identifying miRNA-mediated cleavage on a transcriptome-wide scale. However, current efforts have not yet detected miRNA-mediated cleavage of circular RNAs using conventional degradome sequencing, largely due to the absence of 3◻ poly(A) tail on circular RNAs. To overcome this limitation, we modified library preparation for degradome sequencing to enrich the decaying transcripts of circular RNA without poly(A) tails (Figure 5A) and developed a computational pipeline to identify the cleavage sites in poly(A)− RNA (Figure S7C). The enrichment and accurate identification of cleavage sites from customized libraries were assessed based on the distribution of degradome reads (Figure 5B, 5C, 5D and Figure S7B) and cleavage transcripts spanning the back-splicing sites (Figure 5E, 5F, 5H, and Figure 6D). Ultimately, 12 circular RNAs were identified as cleavage transcripts mediated by miRNAs (Figure 5H). Overexpressing miR166 downregulated the expression of *circ-NHLRC2* (Figure 5I and 5J), which provides evidence for miRNA-mediated cleavage of circular RNA. However, further investigation is needed to elucidate similarities and differences in miRNA-mediated cleavage mechanisms between circular RNAs and linear transcripts, the latter of which involves the major effector endonuclease AGO1 [47].

Quantitative analyses allowed us to identify core circular RNAs involved in the response to GA and NAA. A total of 1,390 circular RNAs modulated by both GA and NAA could function as potential mediators of cross-talk between GA and NAA (Figure 6C). Further investigation of the biological functions of these hormone-induced circular transcripts indicated that these circular RNAs might be involved in processes related to plant hormones, second messengers, cell inclusion bodies, and plant organs (Figure S8). Notably, genes related to rapid growth, particularly those affecting the cell wall, cellulose, and lignin, also generated significantly differentially expressed circular RNAs, which was consistent with the growth-related phenotypes of the hormone-treated seedlings (Figure 6E). In addition, circular RNAs derived from 15 GA-related and 87 NAA-related genes, were likely to mediate biogenesis, function and transport processes of gibberellin and auxin molecules (Figure S8B and S8C). Together, these findings indicate a link between circular RNAs and hormones, although further experiments such as the transgenic expression of circular RNAs associated with hormone regulation are required to elucidate the underlying mechanisms in bamboo.

In previous studies, genes from moso bamboo were overexpressed in *Arabidopsis* and *Oryza sativa*. In total, 55 genes from previous studies have been reported to play key roles based on Agrobacterium-mediated transformation [48–60]. Among these overexpressed genes, we found 12 genes to generate circular RNAs in this study (Table S8). For example, *CWINV4*, encoding cell wall invertase, increased plant height and dry weight in *Arabidopsis* [60] and generated two circular RNAs. It will be interesting to investigate the biological role of these circRNAs in bamboo. Here, we overexpressed 6 candidate circRNAs in rice and observed influences on plant height phenotypes from *cir-AGO1A* and *cir-GID1* (Figure 6), which suggested that some circRNAs have biological roles. The precise roles of these circRNAs could be investigated in the endogenous bamboo system if Agrobacterium-mediated transformation efficiency improves in moso bamboo in the future.

Overall, our study not only enhances the view of hormone responses in moso bamboo (**Figure 7**, module I) but also expands the understanding of circular RNAs, including biogenesis (Figure 7, module II), miRNA-mediated degradation (Figure 7, module III), and function (Figure 7, module IV). In particular, our results dissected the interplay between hormones and circRNA metabolism (Figure 7 and Table S9). Firstly, the expression levels of circular RNAs exhibited high correlation with those of 96 splicing factor genes, including 5 GA-induced factors and 2 NAA-induced factors (Figure 7 and Table S9), which was consistent with our findings that alteration of circRNA levels upon GA treatment was more sensitive than that upon NAA treatment (Figure 7, module II: circRNA biogenesis). Secondly, circular RNAs were cleaved by RISC, including 3 GA-induced RISC and 1 NAA-induced RISC. We further found 7 circRNAs that were subject to miRNA-mediated cleavage in response to hormone treatment, suggesting that hormones might affect the degradation of circular RNAs by modulating the expression of RSCs (Figure 7, module III: circRNA degradation). The steady-state abundance of circRNAs will depend on the balance between these processes of circular RNA biogenesis and degradation. Finally, we revealed potential functions of circRNAs, including regulation of AS, translation, and gene expression (Figure 7, module IV: circRNA function). In total, 379 GA-induced circRNAs and 223 NAA-induced circRNAs were positive correlated with AS of their host gene, indicating that these differentially expressed circular RNAs might alter AS events in response to hormone signals. Notably, several circular RNAs, such as *circ-SF3B2-1*, *circ-EMA1*, and *circ-MYBH*, might be involved in the metabolism of both circular RNAs and hormones by affecting splicing of their linear mRNA counterparts (Figure 7).

**Figure 7.**
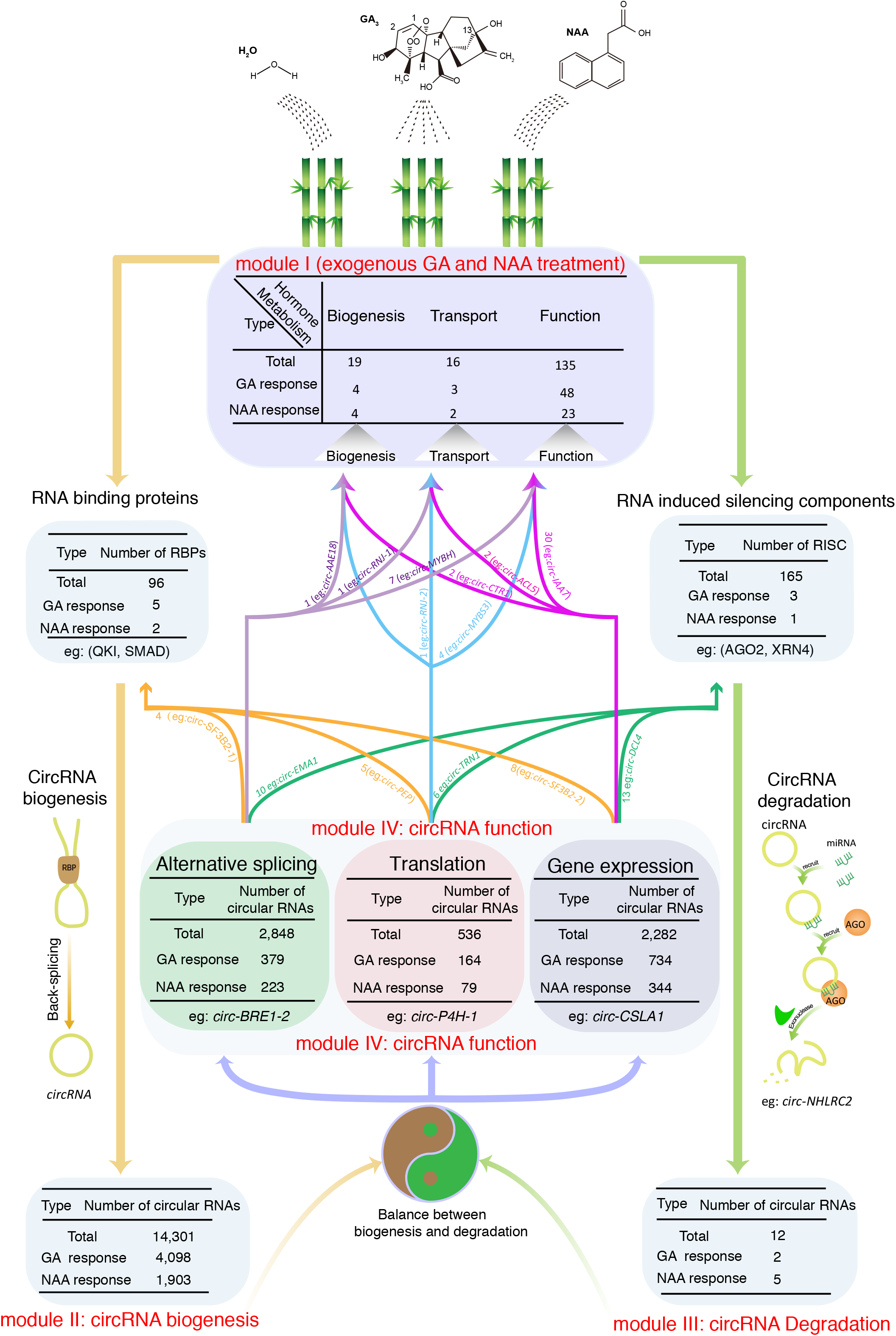
The potential interplay between hormone and circRNA metabolism. Module I: circular RNAs originated from hormone-related genes exhibited dynamic expression upon exposure to GA and NAA. Module II: hormones could affect circRNA biogenesis by regulating several splicing factors. Module III: hormones could regulate the degradation of circular RNAs though modulating RSC. Module VI: Functional circRNAs might regulate hormone metabolism by regulating splicing and transcription of their linear cognates or generating potential proteins from translatable circRNAs.

We further provide evidence that translatable circRNAs might generate functional peptides. We found that translatable circRNAs exhibited dynamic expression upon exposure to hormones. For example, *circ-MYBS* generated a detectable unique peptide, which might play potential roles in regulating hormone metabolism. Moreover, circRNAs could regulate transcription of their parental genes (Figure 7, module IV: circRNA function). A striking example was *circ-DCL4*, originated from miRNA-related genes, which tended to reduce the accumulation level of its linear RNA. The expression of genes related with hormone metabolism might be regulated by several circRNAs, including circ-*CTR1*, circ-*ACL5*, circ-*IAA7 et al* (Figure 7). Taken together, our results highlight the potential interplay between hormone metabolism and circular RNA biogenesis, function, and degradation.

## Material and methods

### Plant materials and hormone treatments

After treatment with GA (100 μM), NAA (5 μM) and ddH_2_O for 4 h, 4-week-old whole seedlings of moso bamboo were collected and transferred to liquid nitrogen. Total RNAs were isolated with the RNAprep Pure Plant Kit (Catalog No. DP441, Tiangen, Beijing, China), and concentrations and quality were determined using 2% agarose gel analysis and NanoDrop quantification before libraries were constructed and high-throughput sequencing performed.

### Library preparation and sequencing for circular RNAs and linear RNAs

To ensure consistency and comparability of different omics, total RNA from each biological repeat were divided into two tubes for circular and linear RNA sequencing. For circular RNAs, ribosomal RNAs were depleted using the Ribo-Zero™ Magnetic Kit (MRZPL1224) and then the samples were incubated with 3U RNase R for 15 min at 37°C. The other group for conventional RNA-seq was incubated with oligo(T) magnetic beads to enrich poly(A)+ RNAs. Subsequently, these RNAs were used for library preparation and subjected to sequencing with Illumina HiSeq 2000.

### Bioinformatics analysis of circular RNAs

For each sample, FASTQ reads were firstly filtered using the HTQC package (v-1.92.1) [61] to remove low-quality reads using default parameters. Clean reads were then mapped to the reference genome [62] using TopHat (v-2.0.11) [63]. Subsequently, the remaining unmapped reads were aligned to the genome using the TopHat-Fusion algorithm (v-2.0.11) for identifying circular RNAs with the CIRCexplorer2 annotation program [7]. Moreover, AS events were detected using CIRCexplorer2 *denovo* [7]. Mutually inclusive circular RNAs were detected by overlapping exons following a previous study [24]. For evolutionary analysis, circular RNAs in *Arabidopsis* and *Oryza sativa* were also identified by the CIRCexplorer2 annotate program with the same parameters as moso bamboo using publicly available data [28, 29]. Reference sequences and annotation of *Arabidopsis* (TAIR10) and *Oryza sativa* (MSU6.1) were obtained from the TAIR and the MSU Genome Annotation Project Database, respectively. We used the bamboo circular RNAs as queries against those of *Arabidopsis* and *Oryza sativa* using BLASTN with –evalue 0.01 and resulting hits sharing below 50% similar stretches of sequence were removed.

To compare circular RNA expression among different samples, we first normalized circular RNA abundance with RPM based on reads spanning the back-spliced junction to the total number of mapped reads (units in million). *P*-values and False discovery rates (FDRs) were calculated using the DEGseq package [64]. Differentially expressed circular RNAs were detected with fold change >= 2, *P*-value < 0.01 and FDR <= 0.01 as cutoff. PCC was calculated by cor() function in R package.

### Experimental validation of circular RNA

For RNase R treatment and PCR amplification validation we used the protocols from our previous study [65]. Briefly, 10×RNase R Reaction Buffer, RNase R and DEPC-treated water was added and incubated for 15 min at 37 °C for RNase R digestion to take place before adding phenol–chloroform–isoamyl alcohol to stop the exonuclease digestion. The sample was then centrifuged at 13,000 g at 4 °C for 5 min and the supernatant was transferred to a new 1.5-ml RNase-free tube containing the following salts: LiCl, glycogen and pre-chilled absolute ethanol (−20°C) and finally inverted gently and stored at −80°C for 1 h. Subsequently, RNase R-treated and control samples were reverse-transcribed to form cDNA for circular RNA amplification with 40 cycles using divergent primers. PCR products with predicted sizes were dissected from an 2% agarose gel and directly sequenced. Divergent primers and convergent primers were designed to detect the candidate circular RNA and positive control for linear RNA, of which both are listed in **Table S10**.

### Functional annotation of moso bamboo

Core spliceosomal factors, RNA-binding proteins including splicing factors, miRNA-related genes, translation-related genes, fast growth–related genes, and hormone-related genes were annotated using BLAST2GO [66] with default options. GO enrichment analysis was performed by BiNGO [67]. The phylogenetic tree of QKI was prepared with ClustalX1.83 and iTQL [68]. The KH domain of QKI was detected by CDD [69] and visualized by GSDS 2.0 [70].

### Protoplast isolation, plasmid construction, and transfection

Protoplasts of moso bamboo were isolated and transformed as previously described [71]. Briefly, shoots of seedlings were immediately transferred into a culture dish containing enzymatic solution and digested for 3 h at 25°C with gentle shaking (50 rpm) in the dark. After adding CPW11M of an equal volume consisting of cell and protoplast washing (CPW) solution at PH5.7 mannitol, protoplast pellets were obtained from the miscible solution filtered through two layers of medical gauze and centrifuged for 3 min at 1200 rpm to remove the supernatant. After an additional two washing steps using the CPW11M solution, the protoplast pellets were resuspended at a concentration of 10^5^~10^6^ protoplasts in 1 mL using MMG solution.

In this study, pUC22-35s-sGFP was used as the backbone for the construction of the overexpression and RNA interference (RNAi) vectors. To overexpress *circ-DCL4*, *circ-PKL*, *circ-CSLA1*, *circ-BRE1-1*/*circ-BRE1-2* and the third linear exon of *NRT1*, the endogenous exons were cloned and flanked by 139-bp inverted complementary flanking introns using overlapping PCR [72]. The two inverted complementary exons (271 bp) from *PedQKI* (PH02Gene29161) were flanked across the intron of *Cunninghamia lanceolata* to construct the vector for *PedQKI* RNAi. *circ-NHLRC2* and miR166 were amplified and flanked by the promoter (CaMV 35S) and NOS terminator using overlapping PCR. Then, two resulting products were inserted in the plasmid to enable the co-expression of both *circ-NHLRC2* and miR166.

The recombinant plasmids were transfected using the PEG-mediated method. For each sample, 100 μL plasmid, protoplast and 110 μL PEG solution were mixed and incubated at 25 °C. After incubation, the mixture was added with 5 mL CPW11M solution and centrifuged for 3 min at 1200 rpm. The remaining protoplasts without supernatant were resuspended with 10 mL CPW11M and centrifuged again at 1200 rpm for 3 min. Finally, after adding 10 mL CPW11M solution, the transfected protoplasts were incubated at 25 °C in the dark for 12–20 h. The expression of transcripts including circular RNA, mRNA and pre-miRNA was detected by semi-quantitative RT-PCR using divergent or convergent primers **(Table S10)**.

### Identification of alternative splicing events from RNA-seq

Low-quality reads were cleaned with default parameters using the HTQC package (v-1.92.1) [61], and the remaining reads were mapped to the genome [62] by TopHat (v-2.0.11) [63]. Aligned reads were used to assemble transcriptome annotation applying Cufflinks (v-2.1.1) with default parameters [73]. Differential expression alternative splicing (AS) events were detected by rMATS.3.2.2 with the following option “-a 8 -c 0.0001 - analysis U” [74]. The parameters for -a and -c represented anchor length and the cutoff splicing difference, respectively. The default anchor length was 8 for RNA-seq splice-aware alignment. The default cutoff splicing difference was 0.0001 for 0.01% difference. To determine whether the frequency of AS events located in transcribed region of circular RNAs was significantly higher than other region of linear RNA, we randomly selected the same number transcribed regions without generating circular RNAs and the number of sequences affected by AS was calculated. After repeating the process 1000 times, we determined the mean value and the standard deviation of the simulation data. PCCs between circRNAs and four types of AS events were calculated using the expression profiles of circRNAs and AS events, which was represented by back-splicing junction reads for circRNAs and splicing junction for AS events, respectively.

### Construction of customized degradome sequencing libraries

Considering that the decay of circular RNAs lacking 3’ poly(A) tail of circular RNAs cannot be detected by conventional degradome sequencing. A modified degradome library preparation was developed based on 5’ RACE library preparation. The Dynabeads™ mRNA Purification Kit was used according to the manufacturer’s instructions to extract total RNAs from 4-week-old bamboo seedlings treated with GA, NAA, and H_2_O. Total RNAs were then separated into poly(A)− and poly(A)+ groups. Subsequently, rRNAs were removed from the poly(A)− group using the Ribo-Zero™ Magnetic Kit. The free 5’ monophosphate of the cleavage transcripts of poly(A)− and poly(A)+ RNA were both ligated to 5’ RNA adapter. Following reverse transcription with Biotinylated Random Primers, the cDNA library was sequenced using Illumina Hiseq2000.

### Bioinformatics analysis for customized degradome sequencing

For customized degradome sequencing, we developed a computational strategy to detect and compare the accumulated decay events termed as degradome peaks with candidate miRNA-mediated cleavage sites in circular RNAs. Firstly, degradome sequencing reads were aligned to the genome using Bowtie2 (v2.2.1) with default parameters [37]. The mapping reads were calculated at the 5’ end alignment positions and counted as 1/n for counting the distribution of reads and the degradome peaks, in which n is total number of mapped reads. The degradome peak was selected by the following cutoff: the percent abundance of cleavage sites was 50% or more in each continuous 21 nucleotides consisting of the 10 nucleotides upstream and downstream of cleavage sites. The *P*-value cutoff of cleavage sites calculated by binomial test was determined as 0.05. Then, the upstream and downstream 25 nucleotides of cleavage sites were aligned to mature miRNAs by RNAplex [38] to identify the target RNA, in which the minimum free energy ratio (MFEratio) was set as 0.7 or less and the sliced sites were in the ±1 region around cleavage sites. To further detect cleavage sites spanning back-splicing sites, we firstly extracted the upstream and downstream 50 nucleotides of back-splicing sites from circular RNAs including degradome peaks. Then, all reads from poly(A)− library were aligned to back-splicing junction regions of circular RNAs using Bowtie2 with default parameters [37]. Finally, the cleavage of circular RNAs supported by degradome reads was identified.

### Small RNA sequencing and bioinformatics analysis

To rapidly and effectively obtain high-quality small RNAs for mock and hormone-treated samples, PEG8000 precipitation was utilized to isolate the small RNAs from 5 μg of total RNA for each library and then 3’-adapter and 5’-adapter were successively ligated to small RNAs before reverse transcription. Finally, the DNA product was enriched using 3.5% PAGE and bands of approximately 200 bp were isolated before sequencing using HiSeq 2500.

The raw RNA reads were processed to remove 5’ and 3’-adapters using the fastx-clipper function of the FASTX toolkit. The filtered reads were subjected to alignment against the Rfam database using bowtie 2 (v2.2.1) [37], to remove the common RNA families (rRNA, tRNA, snRNA, and snoRNA).The remaining sequences were aligned against miRBase database (v21) [75], using BLAST algorithm to identify miRNAs, allowing for e-value<0.05 and three mismatches in total between targets reads and known miRNAs. Completely matched sequences were deemed as conserved miRNAs and other sequences with no more than three mismatches or gaps were considered as variant miRNAs. Reads per million (RPM) mapped reads were used to normalize the expression of miRNA from different libraries.

### Identification and annotation of ORFs from circular RNAs

To predict and annotate the cORFs, circRNA sequences excluding introns were multiplied four times for ORF prediction **(Figure S5)** applying Transdecoder with lowest length >10aa [76]. Subsequently, three protein databases including non-redundant protein sequences of NCBI (NR) [77], UniProt (https://www.uniprot.org) and PsORF [33] using BLASTP to detect their homologous proteins with known function using following parameters: Identities >80, E-value <0.01 and the length of alignment >50%. By the same measurement, uORFs and dORFs derived from linear RNAs were predicted and annotated. To detect unique peptides for each of the ORFs in cORFs, uORFs, dORFs, and pORFs, we obtained raw proteome data based on label-free and tandem mass tags (TMT) approaches [78, 79]. We then performed four independent searches to identify the peptide matching ORFs using MaxQuant with standard parameters [80]. Subsequently, we removed the peptides matching more than one ORF, and the ORFs with at least one unique peptide were retained.

### Transformation procedure for six circRNAs

The pCAMBIA1390 vector including six circRNAs and flanking inverted complementary intron sequences were transformed into Kitaake (*Oryza sativa* ssp *japonica*). The mature embryos were used as the material for callus induction and Agrobacterium-mediated transformation. The hygromycin-resistant callus was transferred into differentiation medium for regeneration. To validate transformation, genomic DNA from rice leaves was extracted using CTAB methods, and the expected fragments were amplified using primers of hygromycin genes. Expression levels of the six circRNA transcripts were further detected by RT-PCR using divergent primers (Table S10).

## Data availability

Raw sequencing data from mRNA sequencing, circular RNA sequencing, degradome sequencing, and small RNA sequencing are available under accession numbers CRA007877 in NGDC (https://ngdc.cncb.ac.cn/gsa). The codes have been submitted to BioCode under accession number BT007322.

## CRediT author statement

**Yongsheng Wang**: Methodology, Investigation, Software, Validation, Writing - original draft. **Huihui Wang**: Investigation, Validation. **Huiyuan Wang**: Software, Data Curation. **Ruifan Zhou**: Investigation, Validation. **Ji Wu**: Validation. **Zekun Zhang**: Software. **Yandong Jin**: Validation. **Tao Li**: Validation. **Markus V. Kohnen**: Software. **Xuqing Liu**: Validation. **Wentao Wei**: Validation. **Kai Chen**: Validation. **Yubang Gao**: Software. **Jiazhi Ding**: Validation. **Hangxiao Zhang**: Software. **Bo Liu**: Conceptualization. **Chentao Lin**: Conceptualization. **Lianfeng Gu**: Conceptualization, Writing - review & editing, Supervision.

## Acknowledgments

This work was supported by the National Natural Science Foundation of China Grant (31971734 and 31800566), the National Key Research and Development Program of China (2018YFD0600101), Distinguished Young Scholar Program of Fujian Agriculture and Forestry University (xjq202017), the Scientific Research Foundation of Graduate School of Fujian Agriculture and Forestry University (324-1122yb061), and the Forestry Peak Discipline Construction Project from Fujian Agriculture and Forestry University.

## Supplementary materials

**Figure S1 GO enrichment analysis for the homologous circular RNAs detected in three plants.**

GO terms in the *x*-axis are categorized into three types with different colors. The *y*-axis indicates the percentage (left *y*-axis) and number (right *y*-axis) of host genes including homologous circular RNAs.

**Figure S2 Length of flanking introns.**

Cumulative curve and boxplots of the length of upstream and downstream flanking introns of circRNAs in comparison with control introns.

**Figure S3 Distribution of Pearson correlation coefficients between circular RNAs and RBPs.**

**A**. Bar plot of the distribution of Pearson correlation coefficients between circular RNAs and 24 core spliceosomal components, 92 splicing factors, and 1132 other RNA-binding proteins excluding splicing factors and core spliceosomal components in bamboo. **B**. Scatter plot of distribution of PCCs between circular RNAs and seven RPBs (SF3A, SF3B, NF90, DHX9, FUS, HNRNPL, and QKI) in bamboo.

**Figure S4 Evolutionary trees of human QKI and its 34 homologous proteins in moso bamboo.**

KH domains are indicated as yellow bars.

**Figure S5. Translatable circular RNAs in moso bamboo**

**A**. Overview of the identification and annotation of the potential cORFs. **B**. The histogram plot shows the percentage of homologous cORFs, uORFs, and dORFs with known protein databases. **C**. Phylogenetic analysis of the cORF of *circ-GLO5* and six homologous cORFs from other species. **D**. Number of the translatable cORFs, uORFs, dORFs, and pORFs based on proteomics. **E**. The MS spectra of cORFs from *circ-P4H-1*. The a, b and y ion are indicated in red, green, and blue lines, respectively.

**Figure S6 Homology analysis of RNase L**

**A**. Multiple sequence alignment of RNase L in different species. **B**. The histogram plot shows the number of homologous proteins.

**Figure S7 Construnction and analysis of degradome libraries**

**A**. The histogram plot shows the number of miRNAs and top 15 miRNA families. **B**. The boxplot shows the distribution of reads in the 5◻ UTR, CDS and 3◻ UTR from poly(A)+ and poly(A)− degradome libraries. **C**. A computational pipeline for identifying cleavage sites based on two types of degradome libraries. **D**. A computational pipeline for detecting decaying transcripts of circular RNAs spanning back-splicing sites.

**Figure S8. Potential function of hormone-induced circular RNAs**

**A**. Heatmap showing expression levels of circular transcripts related to biological process GO terms plant hormones, second messenger, cell inclusion bodies, and plant organs. (B) and (C) Red genes indicate host genes that generated circular RNAs. The light-green box indicates biosynthesis, light purple indicates transport, and light yellow indicates signaling. The events in (B) are in the presence of gibberellin and auxin is used in (C). Black arrows indicate positive effects and black dashed lines indicate negative effects.

**Table S1 Statistical data for sequencing and circular RNAs in seedlings of moso bamboo**

**Table S2 Core spliceosomal factors, splicing factors, and other RBPs with higher PCCs**

**Table S3 Alternative splicing events located in transcribed regions of circular RNAs in the three species**

**Table S4 Identied mature miRNAs and miRNA-related gene in moso bamboo**

**Table S5 CircRNAs and linear RNAs decayed by miRNAs**

**Table S6 Translatable circular RNAs and translation-related proteins**

**Table S7 Differentially expressed circRNAs and GA and NAA hormone-related genes**

**Table S8 Genes from bamboo transformed into *Arabidopsis*, *Oryza sativa*, and *Nicotiana tabacum***

**Table S9 Interplay between hormone (GA and NAA) treatment and circular RNA metabolism**

**Table S10 Primers for validation and transformation of circular RNAs**

