## Supplementary figures and images for "Multi-omics of Circular RNAs and Their Responses to Hormones in Moso Bamboo (*Phyllostachys edulis*)"

### Figure S1

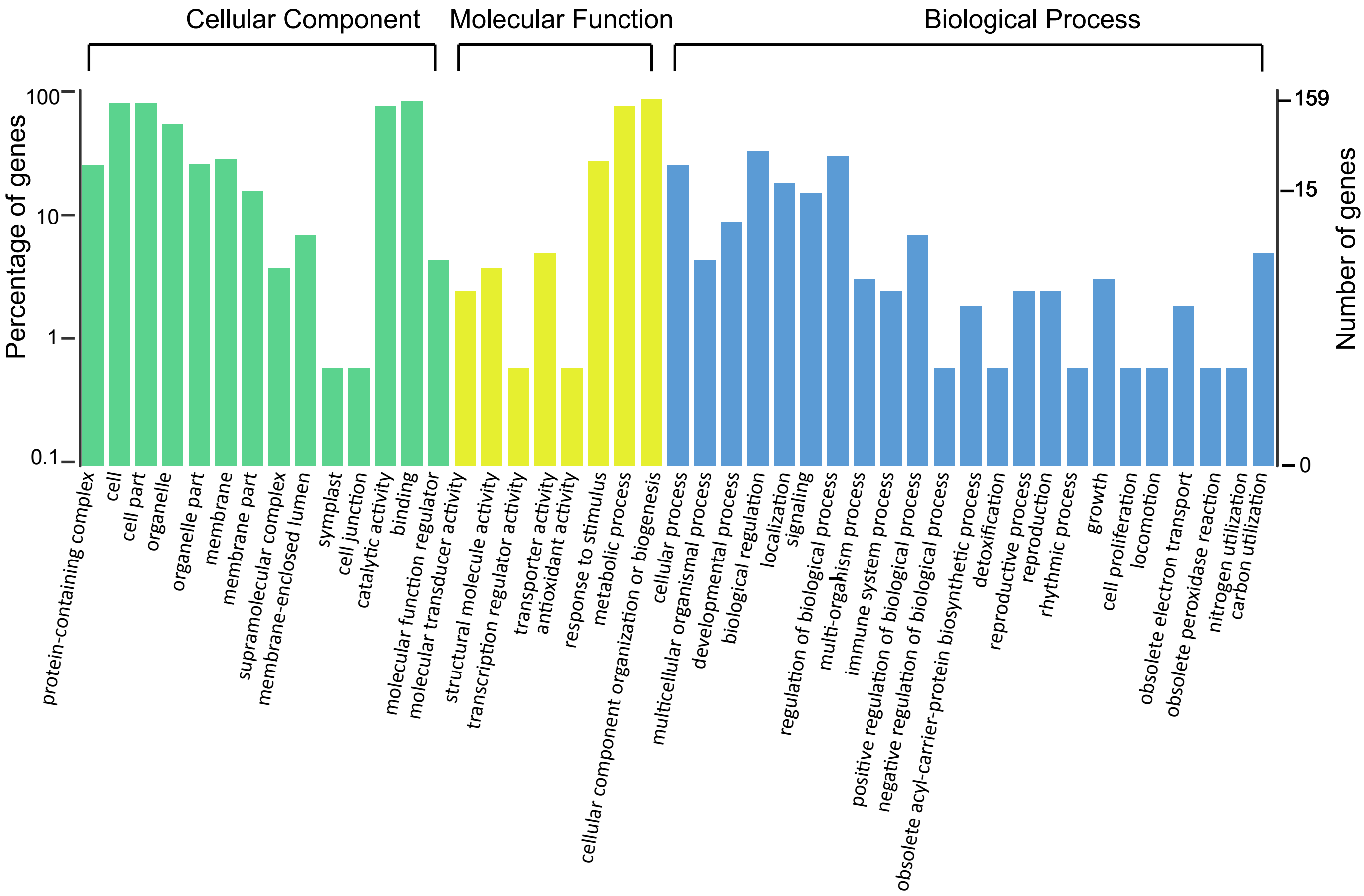

### Figure S2

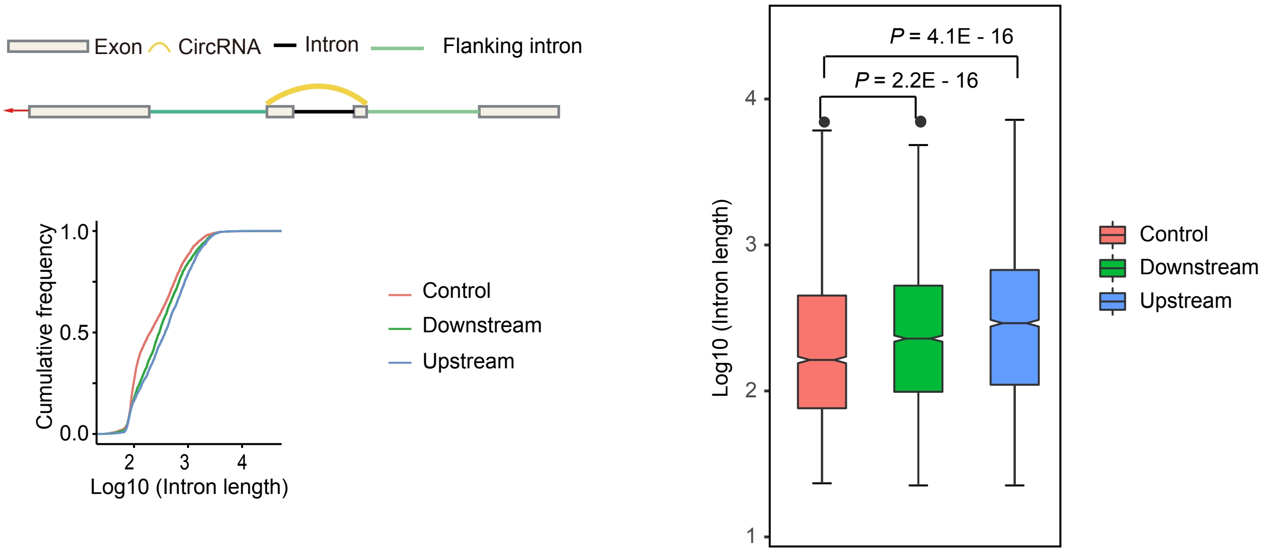

### Figure S3

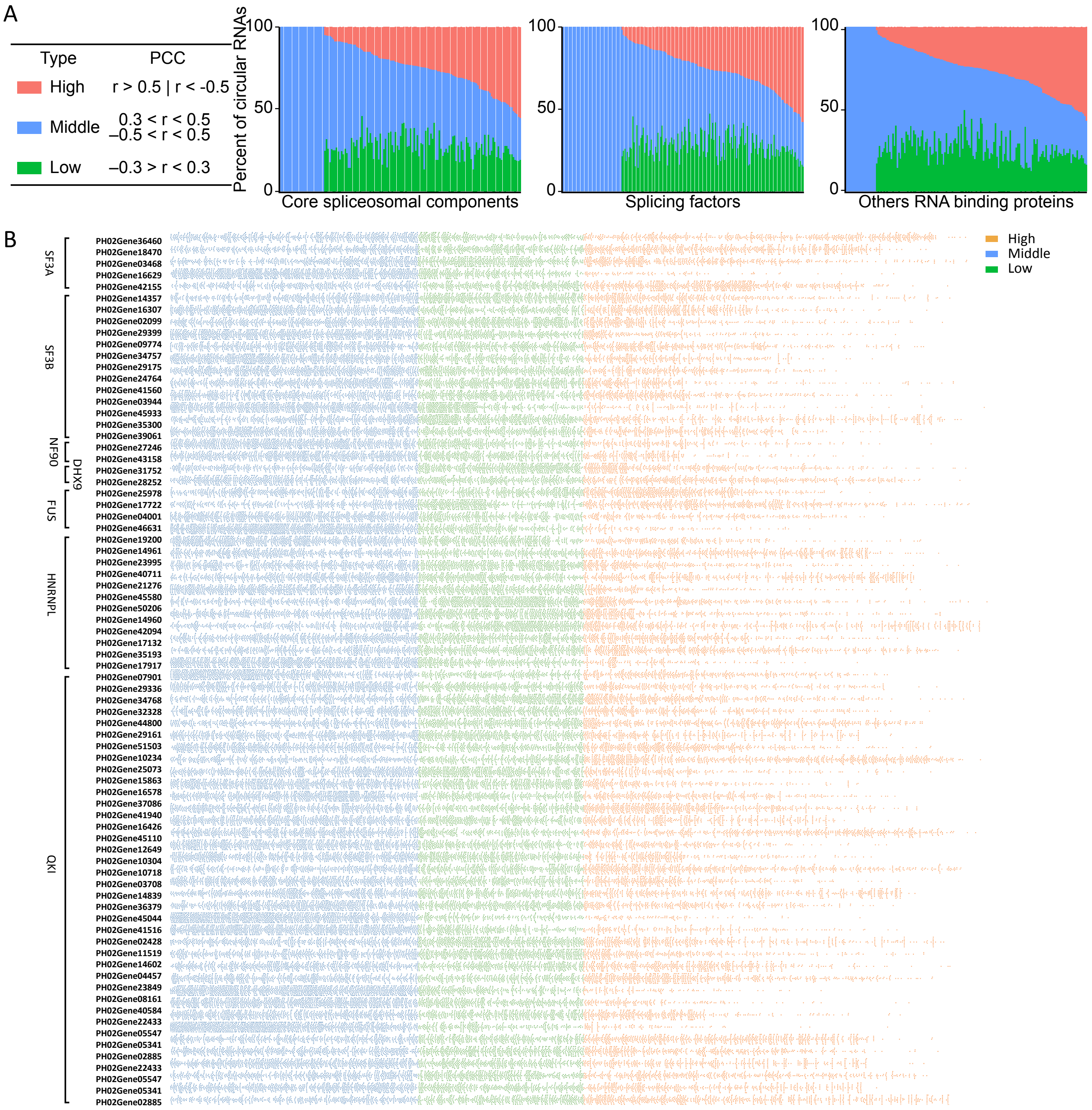

### Figure S4

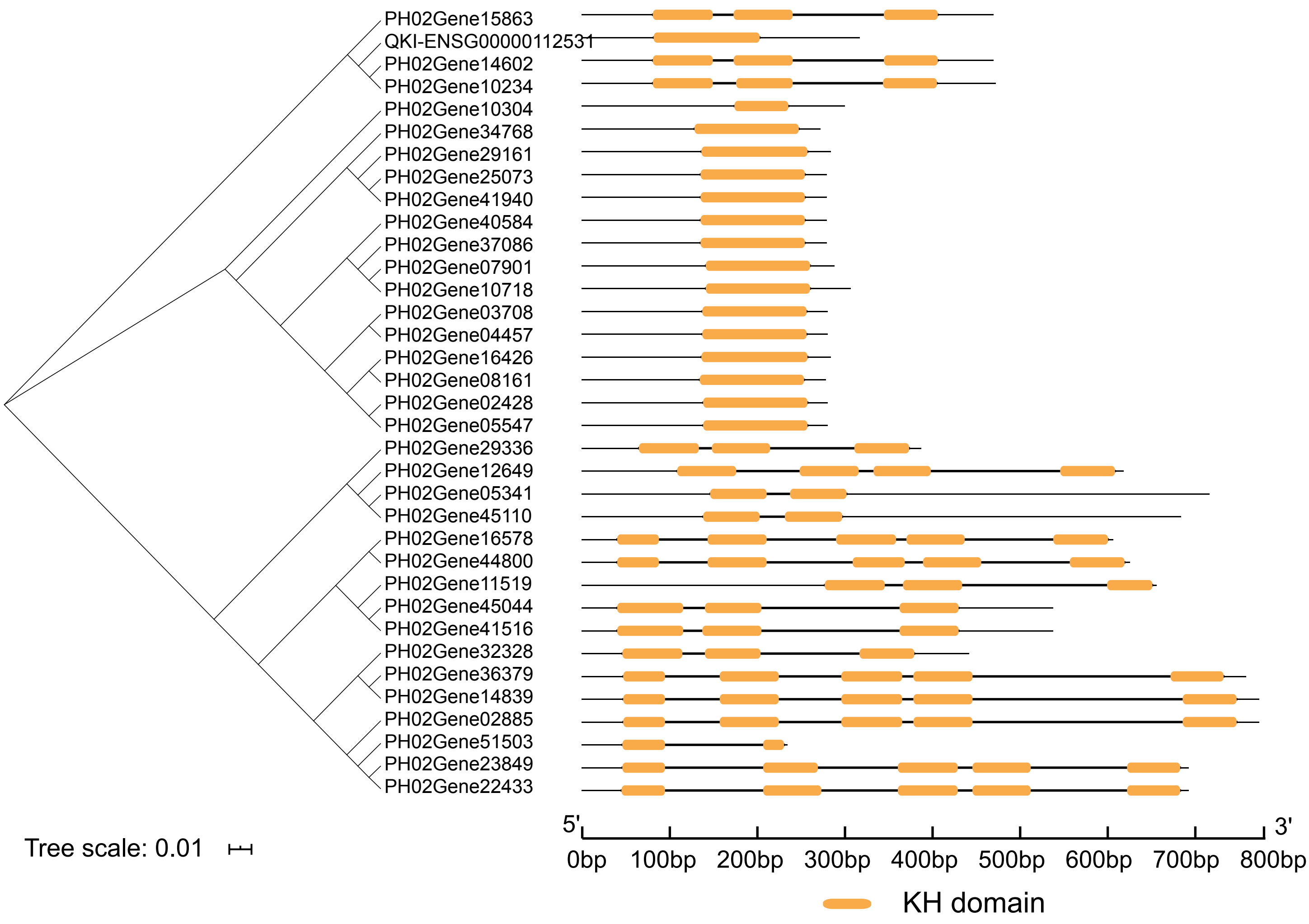

### Figure S5

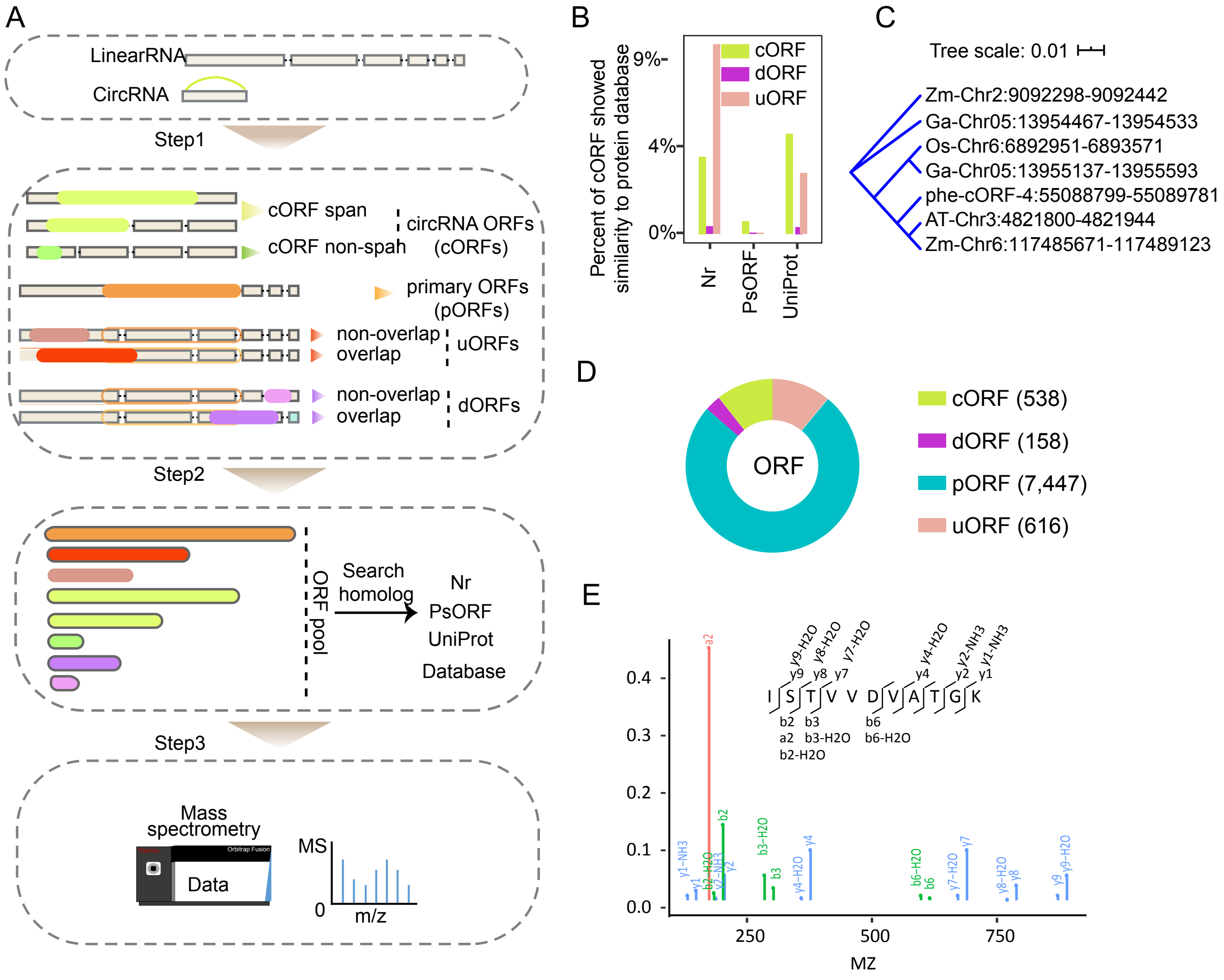

### Figure S6

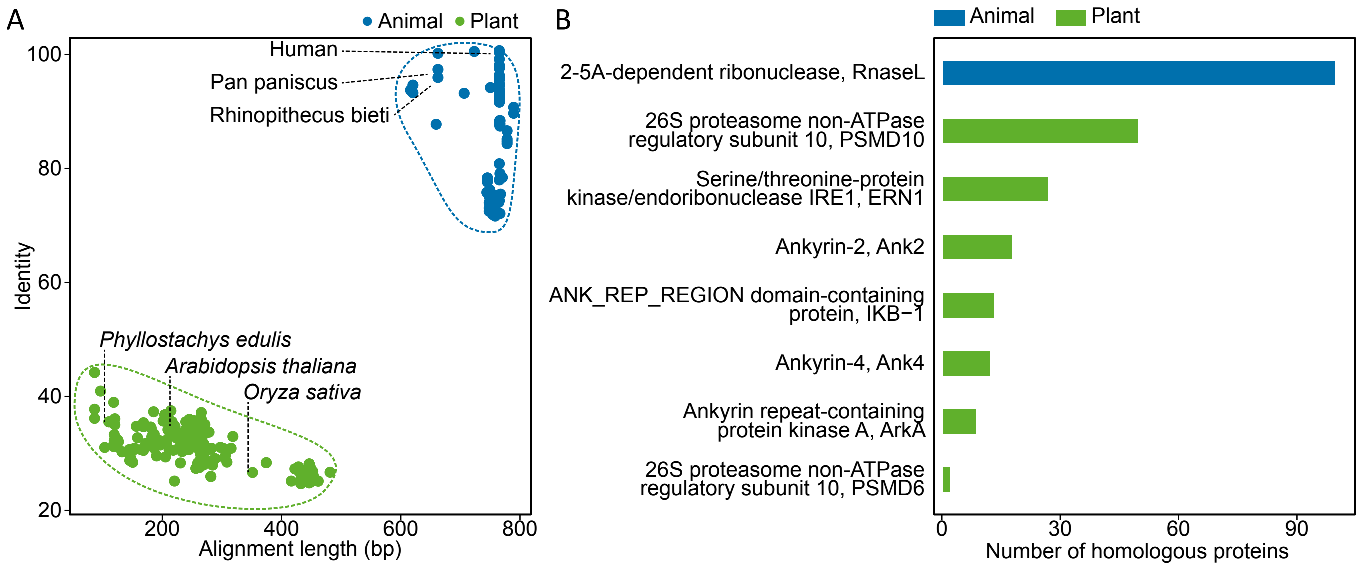

### Figure S7

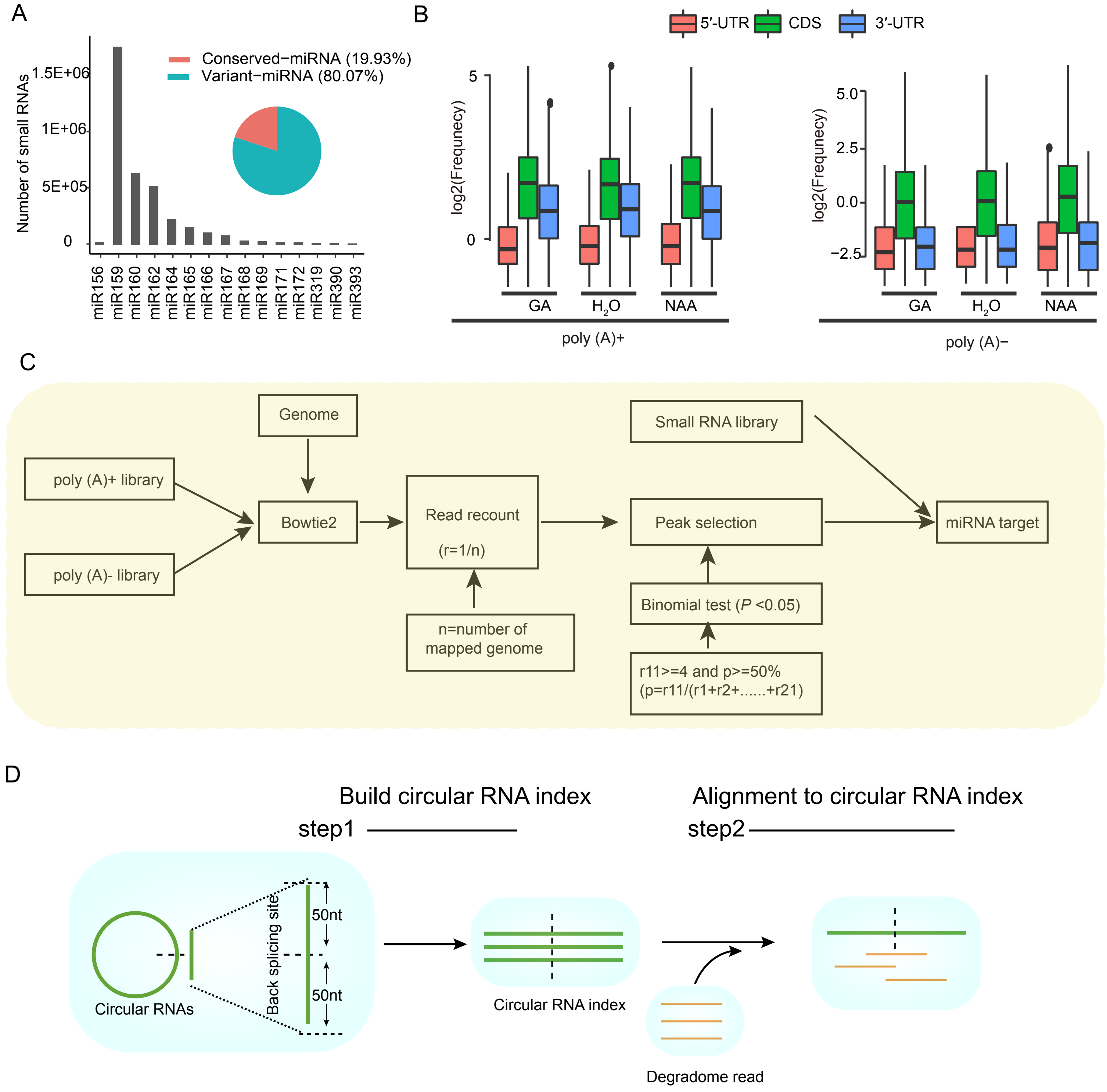

### Figure S8

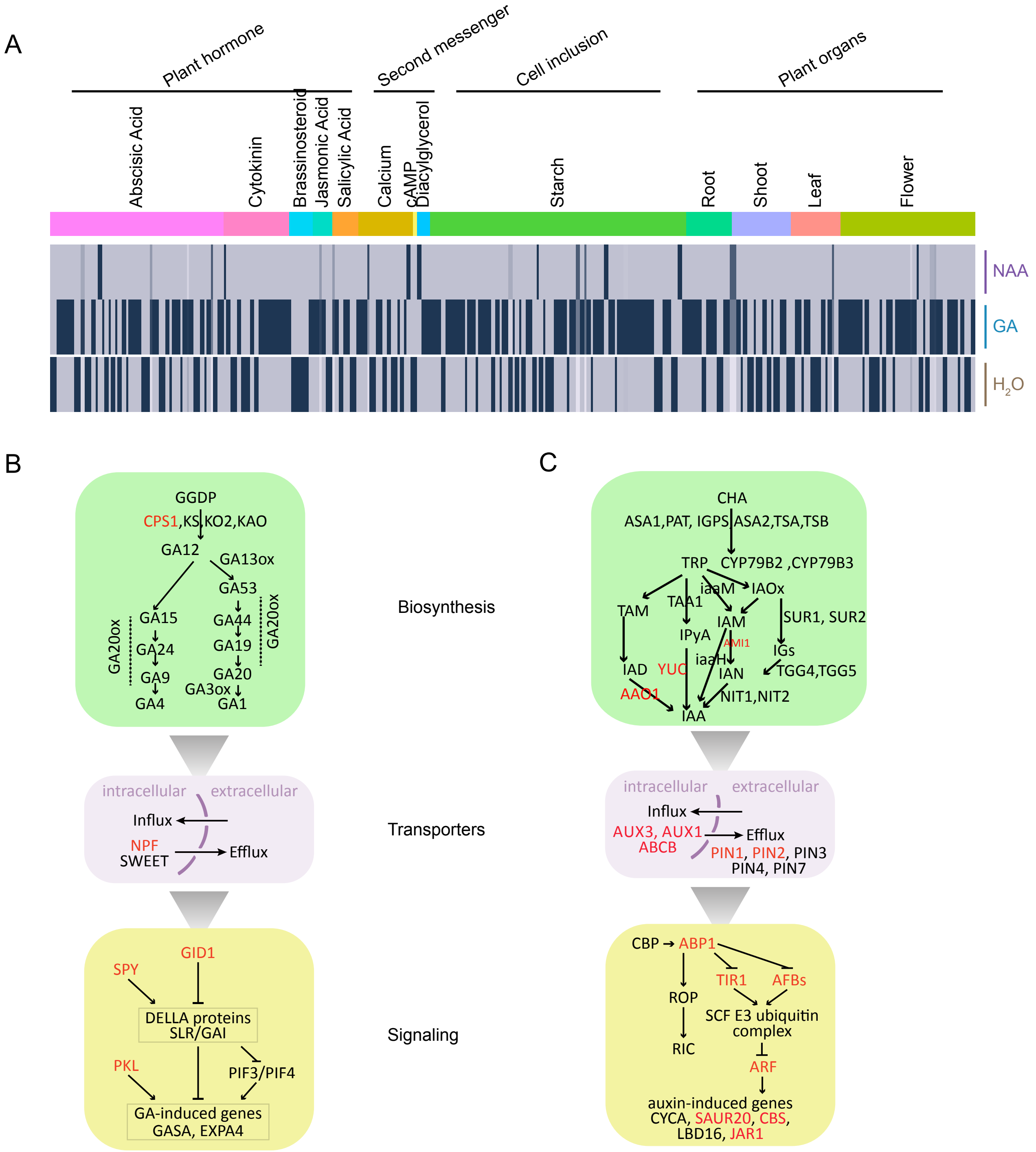

### Figure S9

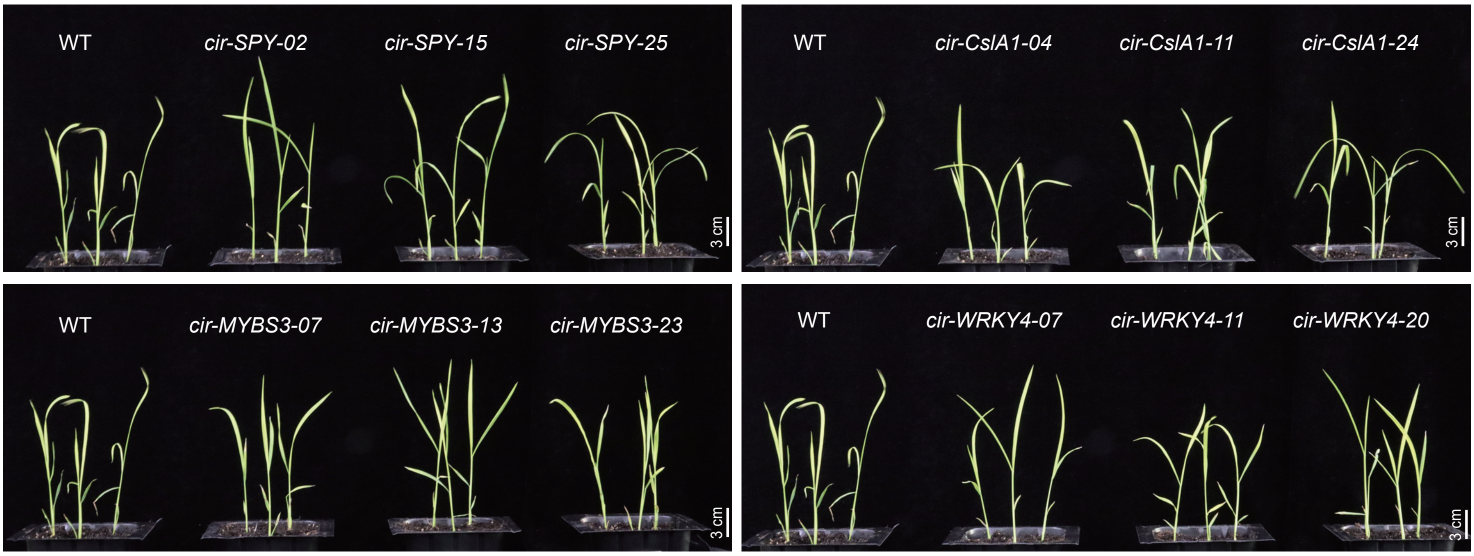
